# Insights into cannabinoid biosynthesis in *Chlamydomonas reinhardtii* : successes with *NphB* and limitations of *CBDAS* expression

**DOI:** 10.64898/2025.12.09.693288

**Authors:** Serge Basile Nouemssi, Ayoub Bouhadada, Remy Beauchemin, Alexandre Custeau, Sarah-Ève Gelinas, Natacha Merindol, Fatma Meddeb-Mouelhi, Hugo Germain, Isabel Desgagné-Penix

## Abstract

The growing legalization of *Cannabis* has increased demand for cannabinoids (CBs). Currently, pharmaceutically relevant CBs are primarily extracted from *Cannabis*, a process that presents challenges related to yield variability and purity. To overcome these limitations, microbial platforms are being explored for the sustainable production of specific CBs. Compared to conventional microbial hosts, the photosynthetic microalga *Chlamydomonas reinhardtii* offers plant-like post-translational modifications, low-cost cultivation, CO_2_ fixation, and low endotoxin contamination, making it a potential chassis for CB biosynthesis. Here, we expressed a codon-optimized, soluble aromatic prenyltransferase (*NphB*_G286S/Y288A_) from *Streptomyces* and *Cannabis sativa* cannabidiolic acid synthase (*CBDAS*) in the nuclear genome of *C. reinhardtii*; transformants were screened for integration, gene expression, protein accumulation, enzymatic activity, and CB production. Our results provide evidence of functional *NphB* expression in *C. reinhardtii*, with *in vitro* CBGA production reaching up to 633 ± 58 µg/L. Although *CBDAS* transcripts were detected under multiple construct designs, neither protein nor CBDA was accumulated, suggesting limitations in expression, localization, or post-translational processing in *C. reinhardtii*. Our study provides the first demonstration of *in vitro* CBGA biosynthesis in the photosynthetic model microalga *C. reinhardtii*, highlighting its potential as a future platform for CB production. It also highlights key challenges in nuclear expression of plant-derived enzymes, such as CBDAS, emphasizing that improved regulatory control, subcellular targeting, and mRNA processing will be required to achieve full pathway reconstruction in *C. reinhardtii*.

**Highlights:**

- First report of active NphB expression in *Chlamydomonas reinhardtii*
- NphB exhibits measurable *in vitro* activity, producing cannabigerolic acid in algal extracts
- Enzyme localization, construct design, and substrate availability critically influence functionality.

## 1. Introduction

*Cannabis sativa* L. (*Cs*), a plant native to Central Asia with a long history of use, is renowned for its diverse phytochemical profile, particularly its phytocannabinoids (CBs). These specialized metabolites hold significant pharmaceutical and economic interest, including notable psychotropic activities [1–3]. CBs have been extensively studied in preclinical settings for treating various conditions, such as chemotherapy-induced nausea and vomiting, pain, multiple sclerosis-related spasticity, and neurodegenerative disorders, including Parkinson’s disease [4–8].

With the increasing legalization of *Cannabis* and CBs for medical and recreational use, the global legal CB industry is projected to reach $58 billion by 2028, with pharmaceutical CBs accounting for about one-third of total revenue [9]. However, current CB production relies primarily on plant extraction, which poses challenges due to variable metabolite abundance and multiple CB analogs in plant extracts [1, 10, 11]. While chemical synthesis offers an alternative, it remains environmentally unfriendly, inefficient, and limited by the structural complexity of CBs [12, 13].

To address these limitations, researchers have explored heterologous CB biosynthesis in alternative hosts, including insect cells, plants, and microbes (bacteria, yeast, and microalgae) [2, 14–23]. This became feasible following the characterization of enzymes involved in CB biosynthesis [24–28]. CB biosynthesis involves multiple interconnected steps across cellular compartments (**Figure 1**). In *C. sativa*, cannabigerolic acid (CBGA), the central CB intermediate, is synthesized in the plastid by *C. sativa* prenyltransferase (*CsPT* or CBGA synthase), which prenylates olivetolic acid using geranyl diphosphate (GPP). CBGA is then converted in the apoplast by Δ^9^-tetrahydrocannabinolic acid synthase (THCAS) or cannabidiolic acid synthase (CBDAS) to produce THCA and CBDA, respectively, the most abundant CBs [2, 3, 10, 29–34].

**Figure 1.**
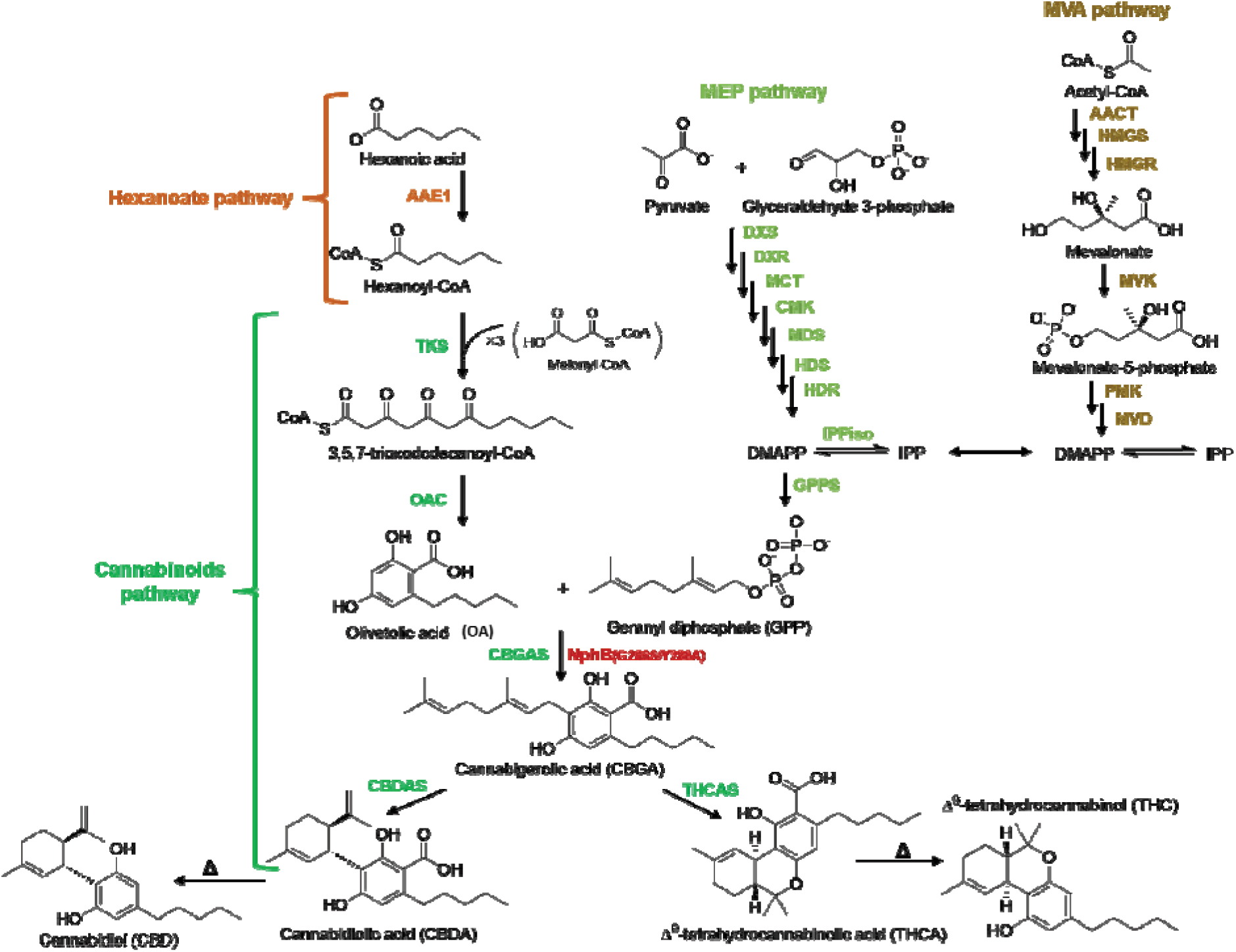
Overview representation of cannabinoids biosynthetic pathway. Represented in red is the double-mutated soluble aromatic prenyltransferase NphB_G286S/Y288A_ from *streptomyces sp*. strain CL190 to substitute the native *Cannabis sativa* cannabigerolic acid synthase (*Cs*CBGAS). Highlighted in orange are enzymes involved in the Hexanoate pathway, the methylerythritol 4-phosphate (MEP) pathway in light green, and the mevalonate (MVA) pathway in brown. The non-enzymatic decarboxylation of final cannabinoids CBDA and THCA is in black. Abbreviations: **AAE1**, Acyl activating enzyme 1; **TKS**, Type III tetraketide synthase; **OAC**, Olivetolic acid synthase; **CBGAS**, Cannabigerolic acid synthase; **CBDAS**, Cannabidiolic acid synthase; **THCAS**, Tetrahydrocannabinolic acid synthase; **DXS**, deoxyxylulose-5-phosphate synthase; **DXR**, 1-deoxy-D-xylulose 5-phosphate reductoisomerase; **MCT**, 4-diphosphocytidyl-2-C-methyl-D-erythritol synthase; **CMK**, 4-diphosphocytidyl-2-C-methyl-D-erythritol kinase; **MDS**, 2-C-methyl-D-erythritol 2,4-cyclodiphosphate synthase; **HDS**, 4-hydroxy-3-methylbut-2-en-1-yl diphosphate synthase; **HDR**, 4-hydroxy-3-methylbut-2-enyl diphosphate reductase, **IPPiso**, isopentenyl diphosphate isomerase, **GPPS**, geranyl diphosphate synthase; **AACT**,acetoacetyl-CoAthiolase; **HMGS**, HMG-CoA synthase; **HMGR**, HMG-CoA reductase; **MVK**, mevalonate-5-kinase; **PMK**, phosphomevalonate kinase; **MVD**, mevalonate diphosphate decarboxylase; **IPP**, isopentenyl diphosphate; **DMAPP**, dimethylallyl-diphosphate. We have adapted this representation from (Degenhardt et al., 2017; Gülck and Møller, 2020; Romero et al., 2020; Zirpel et al., 2017).

Heterologous expression of CBGA biosynthetic enzymes, such as *Cs*PT1 and *Cs*PT4, in yeast and *E. coli* has proven challenging due to their transmembrane plastidial localization, glycosylation requirements, misfolding, and improper compartmentalization [16, 17, 35–38]. Therefore, the bacterial aromatic prenyltransferase NphB from *Streptomyces* sp. strain CL190 has been explored as an alternative CBGA synthase. While wild-type (WT) NphB exhibits low activity and substrate specificity, engineered variants have demonstrated enhanced catalytic efficiency [39–42]. Notably, Valliere et al. identified the *NphB*_G286S/Y288A_ variant, which exhibited an over 100-fold increase in CBGA synthesis, making it a strong candidate for heterologous CB production [43].

Among microbial hosts, the green microalga *Chlamydomonas reinhardtii* presents a promising platform for CB biosynthesis due to its well-characterized genome and closer evolutionary relationship to plants than yeast or bacteria [44]. This unicellular, photosynthetic organism has been widely used for biotechnology applications, including high-value metabolite production [14, 45–47]. Although early studies faced challenges with transgene silencing [48], advances in promoter design, intron incorporation, strain optimization, and chemical treatments have significantly improved heterologous gene expression [48–53]. Furthermore, *C. reinhardtii* has been successfully engineered to accumulate high titers of isoprenoids, a key CB precursor, without compromising native pigment levels or cell fitness [45, 54–56], supporting its potential for CB biosynthesis.

In this study, we establish *C. reinhardtii* as a heterologous host for CB biosynthesis by engineering it to express the *NphB*_G286S/Y288A_ variant (hereafter referred to as NphB) and *CBDAS*. We assessed transgene integration, expression, enzymatic activity, and CBGA/CBDA production under different promoter configurations, genetic constructs, and strains.

We designed a bicistronic construct (C1), expressing *NphB* and *CBDAS* under the chimeric HSP70A-RBCS2 promoter (ARp), and two additional constructs: C2 (*NphB* alone under ARp) and C3 (*NphB* alone under the PSAD promoter (PSADp)). Three *C. reinhardtii* WT strains (CC-125, CC-1690, and CC-5415), as well as the high-expression strains UVM4 and UVM11, were used. This is the first demonstration of functional NphB expression in *C. reinhardtii*, achieving CBGA titers of up to 633 ± 58 μg/L *in vitro*. These findings underscore the potential of *C. reinhardtii* as a sustainable platform for CB production and provide insights into the challenges and opportunities of engineering this host for cannabinoid biosynthesis.

## 2. Materials and methods

### 2.1 Cultivation conditions of microalgal and bacterial strains

Wild-type *C. reinhardtii* strains CC-125 (mt^+^,137c), CC-1690 (mt^+^, Sager 21 gr), CC-5415 (nit1 agg1 mt^+^, Witman g1), and the expression strains UVM4 and UVM 11, were used in this study.

Strains CC-125 and CC-1690 were obtained from the *Chlamydomonas* Resource Center (The University of Minnesota, USA). UVM4 and UVM11 were provided by Prof. Ralph Bock (Max Planck Institute of Molecular Plant Physiology, Germany), and strain CC-5415 was kindly provided by Prof. Martin Jonikas (Jonikas Lab, Princeton University, New Jersey). The microalgal strains were maintained under photo-mixotrophic conditions on Tris-acetate-phosphate (TAP) agar plates [57–59] or in a liquid TAP medium. Liquid cultures were grown in 125 mL flasks on a rotary shaker (120 rpm) under white fluorescent light (50–90 µE m ² s ¹) with a 16 h light/8 h dark cycle at 25 ± 0.5°C.

The bacterial strains *E. coli* DH5α (Invitrogen) and *E. coli* TOP10 (Thermo Fisher Scientific) were used to propagate plasmids, while *E. coli* Rosetta BL21 (DE3) pLysS (Novagen®) was used for recombinant protein expression. Bacteria were cultured in Luria-Bertani (LB) medium supplemented with appropriate antibiotics (100 mg/L ampicillin, 35 mg/L chloramphenicol) at 37°C under shaking at 200 rpm for liquid culture.

### 2.2 PCR amplification, cloning, and expression vector construction

The plasmids pOpt_mRuby2_*aph*VII, containing the *aminoglycoside 3’-phosphotransferase* gene VII (*aph*VII) for hygromycin B resistance [60] and pSL18_*aph*VIII, carrying the *aminoglycoside 3’-phosphotransferase* gene VIII (*aph*VIII) for paromomycin resistance [61], were obtained from the *Chlamydomonas* Resource Center. These plasmids were used as backbones for transgene integration into the *C. reinhardtii* nuclear genome. The plasmid pMAL-c2x (New England Biolabs) was used for the recombinant protein expression in *E. coli*.[60–62]

The coding sequences of *NphB* (*NphB*_G286S/Y288A_) [43], extended Foot-and-Mouth Disease Virus 2A peptide (2A) [62], and *CBDAS* [63] were codon-optimized for *C. reinhardtii* nuclear expression and synthesized (Bio Basic, Markham, Ontario, Canada).

Transgene insertion into the pOpt_mRuby2_*aph*VII vector backbone was performed using Gibson assembly with the NEBuilder® HiFi DNA Assembly kit (NEB), following the manufacturer’s instructions. The required DNA fragments were amplified using Q5 High-Fidelity DNA Polymerase (NEB) and assembled under the control of the HSP70A-RBCS2 fusion promoter (ARp) and the RBCS2 terminator. This resulted in the recombinant expression vector pOpt_mRuby2_NphB-2A-CBDAS (**Figure S1A**), hereafter referred to as construction 1 (C1).

To generate the recombinant vectors pOpt_mRuby2-NphB (construction 2, C2; **Figure S1B**) and pOpt_mRuby2-CBDAS (Construction 4, C4; **Figure S1D**), the *NphB* and *CBDAS* coding sequences were amplified from C1 using primers specific to the vector backbone. The amplified fragments were gel-purified using the GenepHlow™ Gel/PCR Kit (Geneaid) and ligated using the NEB Kinase, Ligase, and *Dpn*I (KLD) enzyme mix, following the manufacturer’s protocol.

A restriction enzyme-based cloning strategy generated additional recombinant vectors: pSL18-NphB (construction 3, C3), pMAL-c2x-NphB, and pMAL-c2x-NphB-2A-CBDAS.

For expression in *C. reinhardtii*, the NphB coding sequence was cloned into the pSL18 backbone under the control of the *PSAD* promoter and terminator (**Figure S1C**). The *NphB* gene was PCR-amplified from C1 using primers designed to introduce *Nde*I/*Xba*I restriction sites at the 5′ and 3′ ends, respectively (**Table S1**). The PCR product was gel-purified, digested with the corresponding restriction enzymes, and ligated into pre-digested pSL18 using T4 DNA ligase (NEB), following the manufacturer’s protocol.

For expression in *E. coli*, the NphB and NphB-2A-CBDAS sequences were cloned into the pMAL-c2x vector under the control of the tac promoter (**Figure S2 A and B**). NphB was fused to the C-terminus of the maltose-binding protein (MBP) tag, while NphB-2A-CBDAS was inserted downstream of the lac operator, without an MBP fusion. The MBP tag enhances recombinant protein’s expression, stability, and solubility and facilitates purification [64, 65]. Coding sequences were amplified from C1 using primers incorporating *BamH*I/*Hind*III sites (for MBP-NphB fusion) or *Nde*I/*EcoR*I (for NphB-2A-CBDAS). PCR products were gel-extracted, digested, purified, and ligated into the appropriately pre-digested pMAL-c2x vector using T4 DNA ligase (NEB).

Table S1 provides a complete list of primers, with introduced restriction sites indicated in bold and underlined.

### 2.3 Transgene expression in the prokaryotic system, screening, and protein production

Recombinant expression vectors were transformed into chemically competent *E. coli* cells for plasmid propagation. *E. coli* TOP10 was used for C1 after Gibson assembly, while *E. coli* DH5α was used for C2 and C3 following ligation-based cloning. Transformation was performed using the heat shock method according to the manufacturer’s protocol. Transformed cells were plated on LB agar supplemented with 100 mg/L ampicillin and incubated at 37°C for 16 hours. To confirm successful plasmid integration, colony PCR was performed as described by [66]. The reaction was carried out in a 25 µL total volume using Taq DNA polymerase and ThermoPol buffer (NEB) under the following conditions: an initial denaturation at 95°C for 5 min, followed by 30 cycles of 95°C for 30 sec, 54°C for 1 min, and 68°C for 1-3 min, with a final extension at 68°C for 5 min and an infinite hold at 4°C. The plasmids were verified via next-generation sequencing (NGS) at the Massachusetts Institute of Technology (MIT) to confirm sequence integrity.

For recombinant protein production, the purified pMAL-c2x-NphB and pMAL-c2x-NphB-2A-CBDAS vectors were extracted from *E. coli* DH5α and transformed into chemically competent *E. coli* Rosetta BL21 (DE3) pLysS using the heat shock transformation. Transformants were selected on LB agar plates containing 100 mg/L ampicillin and 35 mg/L chloramphenicol and incubated overnight at 37°C. A single PCR-positive colony was picked and grown overnight at 37°C in 12.5 mL LB broth containing antibiotics under shaking at 200 rpm. The overnight culture was inoculated into 250 mL fresh LB broth supplemented with antibiotics and grown at 37°C, 200 rpm, until the OD reached 0.4–0.7. The cultures were then brought to room temperature before induction with 0.5 mM isopropyl-β-D-thiogalactopyranoside (IPTG). Induction was performed at 18°C for 20 hours with shaking at 150 rpm. Following induction, cells were harvested by centrifugation at 17,000 x g and resuspended in the protein solubilization buffer (25 mM Tris HCl, pH 7.5; 150 mM NaCl; 1 mM EDTA, pH 8). Protein extraction was performed by sonication using a Fisherbrand™ Model 505 Sonic Dismembrator (Thermo Fisher Scientific) under the following conditions: total sonication time 8 min; amplitude 41%, pulse on 14 sec; pulse off 35 sec. Cell lysates were centrifuged at 20,000 x g for 30 min at 4°C, and the supernatant containing the soluble protein fraction was collected and stored at -80°C for further analysis.

### 2.4 Nuclear transformation and screening of the microalga *C. reinhardtii*

The nuclear transformation of *C. reinhardtii* wild-type strains was performed by electroporation, following the protocol described by [50, 67], with slight modifications. A total of 2 µg of gel-purified vectors, linearized using *Sca*I, or a combination of *Sca*I and *Kpn*I, was used to transform the microalgae. Electroporation was carried out using the Bio-Rad Gene Pulser Xcell^TM^ system in a 4 mm cuvette under the following conditions: 0.5 kV, 50 µF, 800 Ω. For the UVM4 and UVM11 strains, nuclear transformation was carried out using the glass bead method as described by Kindle [68]. In this case, 2 µg of gel-purified *Sca*I-linearized vector was mixed with 500 µL of *C. reinhardtii* cells resuspended in TAP medium supplemented with 5% polyethylene glycol (PEG). The suspension was vortexed for 15 sec and spread onto selective TAP agar plates.

Transformants were selected on TAP-agar plates supplemented with appropriate antibiotic (hygromycin, 15 mg/L or paromomycin, 15 mg/L) and incubated for 5–7 days until single colonies appeared. Colony enumeration was performed using Open CFU software [69]. Randomly selected colonies were subcultured weekly on fresh TAP-agar plates containing antibiotics for five rounds to ensure stable genomic integration of the gene of interest. Subsequent screening was conducted using high-throughput colony PCR (HT-cPCR) as described by [67].

For the co-expression of C2 and C4 as independent open reading frames (ORFs), both constructs, carrying the same selection marker, were co-transformed in a single procedure. Specifically, 1µg of gel-purified linearized vector from each construct was combined in a 4 mm cuvette containing wild-type *C. reinhardtii* cells, followed by electroporation and screening under the abovementioned conditions.

To verify the integrity of the gene sequences following genomic integration, PCR products from HT-cPCR-confirmed transformants were analyzed by Sanger sequencing at the Centre de Recherche CHU de Québec, Université Laval.

### 2.5 DNA extraction and colony PCR in *C. reinhardtii*

Genomic DNA (gDNA) extraction for HT-cPCR was performed following the protocol described by [67]. Two microliters of gDNA were used for colony PCR, which was carried out using Taq DNA Polymerase with ThermoPol® Buffer under the following conditions: an initial denaturation at 95°C for 5 min, followed by 30 cycles of 95°C for 30 sec, 54°C for 1 min, and 68°C for 1–3 min, with a final extension at 68°C for 5 min and an infinite hold at 4°C. The primer sequences used are listed in **Supplementary Table S1**.

Before characterizing PCR-positive clones for transgene expression, an additional PCR targeting the gene of interest was performed on gDNA extracted from liquid cultures. This extraction followed the protocol described by [70], with slight modifications to assess transgene stability in clones grown for five days in a 25 mL TAP medium. Briefly, 2 mL of algal culture from day five was centrifuged at 17,000 × g for 1 min. The resulting cell pellet was resuspended in 200 µL of extraction buffer (2% cetyltrimethylammonium bromide [CTAB], 100 mM Tris-HCl pH 8.0, 20 mM EDTA pH 8.0, 1.4 M NaCl, and 2% [v/v] freshly added β-mercaptoethanol) by vortexing. Next, 200 µL of chloroform/isoamyl alcohol (24:1) was added, and the mixture was incubated at 65°C for 20 min with shaking at 1400 rpm. After centrifugation at 17,000 x g for 10 min at 10°C, the DNA-containing supernatant was carefully collected by tilting the tube at a 45° angle to avoid contamination from the organic phase and transferred to a new microcentrifuge tube. The gDNA was precipitated by adding 0.7 volumes of isopropanol to the supernatant, followed by a 10 min incubation at room temperature and centrifugation at 17,000 x g for 10 min at 4°C. The pellet was then washed with 1 mL of 70% (v/v) ethanol, air-dried, and resuspended in 50 µL of DNase-free water supplemented with 1 µL of RNase A (10 mg/mL), followed by incubation at 37°C for 15 min. The concentration and purity of the extracted gDNA were assessed by measuring the OD _/_ ratio using a NanoPhotometer® N60/N50 (Implen).

### 2.6 Total RNA extraction and quantitative PCR

Total RNA was extracted from 5-day-old *C. reinhardtii* cultures using the TRIzol reagent (Invitrogen). A 4 mL culture was collected and centrifuged at 12,000 x g for 5 min, after which the cell pellet was resuspended in 500 µL of TRIzol. To enhance cell lysis, samples were flash-frozen in liquid nitrogen, incubated at room temperature for 10 min, and then extracted with 200 µL of chloroform. After centrifugation at 12,000 x g for 10 min at 4°C, the RNA-containing upper phase was carefully transferred to a fresh microcentrifuge tube. RNA was precipitated by adding 500 µL of isopropanol and 100 mM NaCl, incubating for 10 min at room temperature, and centrifuging at 12,000 × g for 10 min at 4°C. The RNA pellet was washed with 1 mL of 75% (v/v) ethanol, centrifuged at 7,500 x g for 5 min at 4°C, air-dried, and resuspended in 50 µL of nuclease-free water. Residual genomic DNA was removed using Turbo DNase (Invitrogen TURBO DNA-free Kit, Thermo Fisher Scientific) following the manufacturer’s instructions. The RNA concentration and purity were assessed by measuring the OD / ratio using a NanoPhotometer® N60/N50.

For quantitative reverse transcription PCR (RT-qPCR), 250 ng of total RNA was subjected to reverse transcription and qPCR amplification in a single reaction using the Luna Universal One-step RT-qPCR Kit (NEB). Reverse transcription was performed at 55°C for 10 min, followed by an initial denaturation at 95°C for 1 min. qPCR cycling conditions included 45 cycles of 95°C for 10 sec (denaturation) and 60°C for 30 sec (extension). A melt curve analysis was conducted from 60°C to 95°C, with an increment of 0.5°C every 5 sec. Primers for *NphB* and *CBDAS* transcripts were designed using the PrimerQuest^TM^ Tool (IDT), while reference gene primers (*histone* 3 and *phosphoglycerate kinase* [*PGK*]) were obtained from [50] (**Table S2**). SYBR Green fluorescence was recorded in the FAM channel of a CFX connect Real-Time System (Bio-Rad), and amplification products were analyzed using Bio-Rad CFX Manager version 3.1. Relative transgene expression levels were determined using the 2^−^ΔΔCt^ method [71]. Each experiment was performed in two technical replicates per clone.

### 2.7 Total protein extraction and western blot analysis

Total protein extraction was performed by sonication as described by [22], with minor modifications. Briefly, *C. reinhardtii* culture (19 mL) grown for five days were harvested by centrifugation at 3,500 x g for 15 min at 4°C. the resulting cell pellets were resuspended in 500 µL of solubilization buffer (51.4 mM Tris-HCl pH 8; 0.75 mM sodium dodecyl sulfate [SDS]; 10 % (v/v) glycerol; 0.02 mM EDTA; 10 mM phenylmethylsulfonyl fluoride [PMSF]; and 2 µL of protease inhibitor cocktail [Sigma-Aldrich]) by gentle pipetting. Cells were lysed by sonication using a Fisherbrand™ Model 505 Sonic Dismembrator (Thermo Fisher Scientific) for six cycles of 3 min at 35% amplitude, with 30 sec pulses on and off. Lysates were centrifuged at 17,000 x g for 30 min at 4°C, and the supernatants, containing the total soluble protein fraction, were flash-frozen in liquid nitrogen and stored at -80°C for subsequent western blot analysis and *in vitro* enzymatic assays. Protein concentrations were determined using the RC DC™ Protein Assay Kit I (Bio-Rad), with bovine serum albumin (BSA) as a standard.

For western blot analysis, 100 µg of total protein was resolved on a 12% (v/v) SDS-PAGE gel at a constant voltage of 100 V for 3 h. Proteins were transferred onto a 0.2 µm polyvinylidene difluoride (PVDF) membrane using the Trans-Blot Turbo Transfer System (Bio-Rad). The membrane was equilibrated in Tris-buffered saline (TBS; 20 mM Tris, 150 mM NaCl, pH 7.6) for 10 min, followed by blocking in 5% (w/v) skim milk prepared in TBS containing 0.1% (v/v) Tween-20 (TBST) for 5 h at room temperature. The membrane was incubated overnight at 4°C in TBST with 5% skim milk containing primary antibodies (1:1000 dilution), either mouse anti-FLAG monoclonal antibody (Millipore Sigma) or mouse anti-HA-tag monoclonal antibody (GenScript). After primary antibody incubation, the membrane was washed three times (10 min each) in TBST and subsequently incubated for 1 h at room temperature in TBST containing 5% skim milk and goat anti-mouse horseradish peroxidase (GAM-HRP) conjugate (1:20,000 dilution). The membrane was washed twice in TBST, and protein detection was performed using Clarity Max Western ECL Substrate (Bio-Rad). Following chemiluminescence detection, the membrane was washed twice in TBST and stained with 0.5% (w/v) Ponceau S solution (0.5% [w/v] Ponceau S in 1% [v/v] acetic acid) for 1 min to assess protein transfer efficiency. Chemiluminescence signals and Ponceau S-stained blots were visualized using the ChemiDoc Imaging System (Bio-Rad) and analyzed with Image Lab software (Bio-Rad). The molecular weight of detected proteins was confirmed using the Precision Plus Protein Dual Color Standards (Bio-Rad).

### 2.8 Enzymatic assays and CBGA detection

*In vitro* enzymatic assays were conducted using three biological replicates in a total reaction volume of 100 µL, following the protocol of [22]. Specifically, 1 mg of soluble total protein extract was added to the reaction buffer (100 mM HEPES, pH 7.5; 25 mM MgCl_2_; 2 mM OA; and 2 mM GPP) and incubated at 30°C for 16 h. Proteins were precipitated with 2% (v/v) trichloroacetic acid to terminate the reaction, and metabolites were extracted using 300 µL of HPLC-grade methanol. The extracts were filtered through 0.22 µm nylon filters, dried using a Savant SPD1010 SpeedVac concentrator (Thermo Scientific), and reconstituted in 100 µL of HPLC-grade methanol. Samples were then diluted 10-fold in the mobile phase (0.1% (v/v) formic acid in Milli-Q water / 0.1% (v/v) formic acid in methanol, 30:70) and analyzed by high-performance liquid chromatography coupled with tandem mass spectrometry (HPLC-MS/MS).

### 2.9 Olivetolic acid supplementation and metabolite extraction

*C. reinhardtii* cells grown for five days in 25 mL of TAP media were supplemented with 2 mM of olivetolic acid (OA). The culture was incubated for 24 h. Cells were harvested by centrifugation at 3,500 x g for 10 min at 4°C. The supernatant was discarded, and the conical tubes were inverted to dry. Metabolites were extracted using methanol at a ratio of 1 mL per 100 mg of fresh biomass. Samples were vortexed for 30 sec and incubated overnight at -20°C. The extract (supernatant) was separated from the pellet by centrifugation at 3,500 × g for 10 minutes at 4°C, followed by sample filtration through 0.22 µm nylon syringe filters. Filtered samples were stored at -20°C for metabolite analysis.

### 2.10 HPLC-DAD and HPLC-MS/MS analysis

To perform the detection of the metabolite by high-performance liquid chromatography (HPLC) with diode-array detection (DAD) and coupled with tandem mass spectrometry (MS/MS), the following cannabinoids and precursors were used as standards: olivetolic acid (OA, CAS 491–72-5) and olivetol (OL, CAS 500–66-3) were purchased from Santa Cruz biotechnologies (Dallas United states). Δ9-tetrahydrocannabinol (THC, CAS 1972-08-3), cannabidiol (CBD, CAS 13956–29-1), cannabinol (CBN, CAS 521–35-7), Δ9-tetrahydrocannabinolic acid (THCA, CAS 23978–85-0), cannabidiolic acid (CBDA, CAS 1244-58-2), cannabigerolic acid (CBGA, CAS 25555–57-1), cannabichromene (CBC, CAS 20675–51-8), cannabigerol (CBG, CAS 25654–31-3), tetrahydrocannabivarin (THCV, CAS 31262–37-0) and cannabidivarin (CBDV, CAS 24274–48-4) were purchased from Agilent Technologies (QC, Canada). Cannabinolic acid (CBNA, CAS 2808-39-1) was purchased from Sigma-Aldrich (ON, Canada).

Initial analyses were performed using high-performance liquid chromatography with a diode array detector (HPLC-DAD). Chromatographic separation of analytes was achieved using an InfinityLab Poroshell 120 EC-C18 column (4.6 × 100 mm, 2.7 mm; Agilent Technologies, QC, Canada) maintained at 30°C. A 10 µL sample was injected into the analytical system. The mobile phase consisted of: (A) 30% of 0.1% (v/v) formic acid in Milli-Q water and (B) 70% of 0.1% (v/v) formic acid in methanol was used. The flow rate was set to 1 mL/min, and the HPLC gradient program was as follows: 0 min, 70% B; 1.0 min, 70 % B; 6.0 min, 77% B; 15.0 min, 90 % B; 15.1 min, 70 % B and 18.0 min, 70 % B. Each run lasted 18.5 min, including column reconditioning before the next injection. The diode array detector was set to acquire the wavelength range of 190 to 400 nm using a deuterium (D_2_) lamp, with UV detection at 220 nm. Compounds were identified by comparing retention time and maximum absorption wavelengths to those of reference standards (**Table S3**, section 2.10). Calibration curves were generated using CBGA standard solutions (10 mg/L and 100 mg/L in HPLC-grade methanol). These solutions were serially diluted to prepare calibration standards at 0.5, 1, 2, 4, 5, 10, 25, 50, and 100 mg/L, each analyzed in triplicate. The standard solutions were injected into the HPLC-DAD system, and calibration curves were plotted based on the area under the curve (AUC) versus analyte concentration. These curves were used to quantify OA, OL, and CBGA in enzymatic and supplementation assays (**Figure S3 A-C**).

Confirmatory analyses were conducted using HPLCMS/MS (Agilent, QC, Canada) equipped with an Agilent Jet Stream ionization source, a binary pump, an autosampler, and a column compartment. Compound separation was achieved using an InfinityLab Poroshell 120 EC-C18 column (4.6 × 100 mm, 2.7 mm). Five microliters of each sample were injected into the column set at 50°C. A gradient method made of (A) 0.1% (v/v) formic acid in Milli-Q water and (B) 0.1% (v/v) formic acid in methanol with a flow rate of 0.5 mL/min was used to achieve chromatographic separation. The HPLC elution program was as follows: 0 min, 70 % B; 7.0 min, 100 % B; 10 min, 100 % B; 12.0 min, 70 % B. The total run time was 14 min per sample. The parameters used in the MS/MS source were set as follows: gas flow rate 8 L/min; gas temperature 220 °C; nebulizer 55 psi; sheath gas flow 12 L/min; sheat gas temperature 380°C; capillary voltage 4500 V; and nozzle voltage 0 V. Agilent MassHunter Data Acquisition (version 1.2) and MassHunter Qualitative Analysis (version 10.0) software were used for data acquisition and processing respectively. Sample analyses were conducted in triggered multiple reaction monitoring (tMRM) acquisition mode, allowing compound identification using authentic standards. MRM transitions and MS/MS parameters used for targeted compounds identification are provided in the supplementary **Table S4**.

### 2.11 Statistical analysis

Statistical analyses were performed using GraphPad Prism (Version 9.4.1, GraphPad Software, US). Data are expressed as means ± standard deviation (SD) from two or three biological replicates, each conducted at least twice in independent experiments.

## 3. Results

### 3.1 Production of NphB and CBDAS enzymes in *E. coli*

Before proceeding with *C. reinhardtii* transformation, we sought to confirm that the codon-optimized *NphB* and *CBDAS* sequences were correctly expressed, translated, and functional in a heterologous system. *E. coli* is a widely used, rapid, and cost-effective model for preliminary expression testing. The two genes *NphB*_G286S/Y288A_ (hereafter *NphB*) from *Streptomyces* sp. strain CL190 [43] and *CBDAS* from *Cannabis sativa* [63] were codon-optimized for expression in the *C. reinhardtii* nuclear genome. Two expression cassettes were designed to evaluate their activity in *E. coli*. The first contained *NphB* alone; the second encoded a fused transgene, *NphB-2A-CBDAS* (**Figure 2A)**. In the latter, both genes were expressed from a single open reading frame (ORF) by linking the *NphB* to the N-terminus of the *CBDAS* using an extended FMDV2A self-cleaving peptide [62]. The construct included N-terminal *2xFlag* and C-terminal *2xHA* tags.

**Figure 2.**
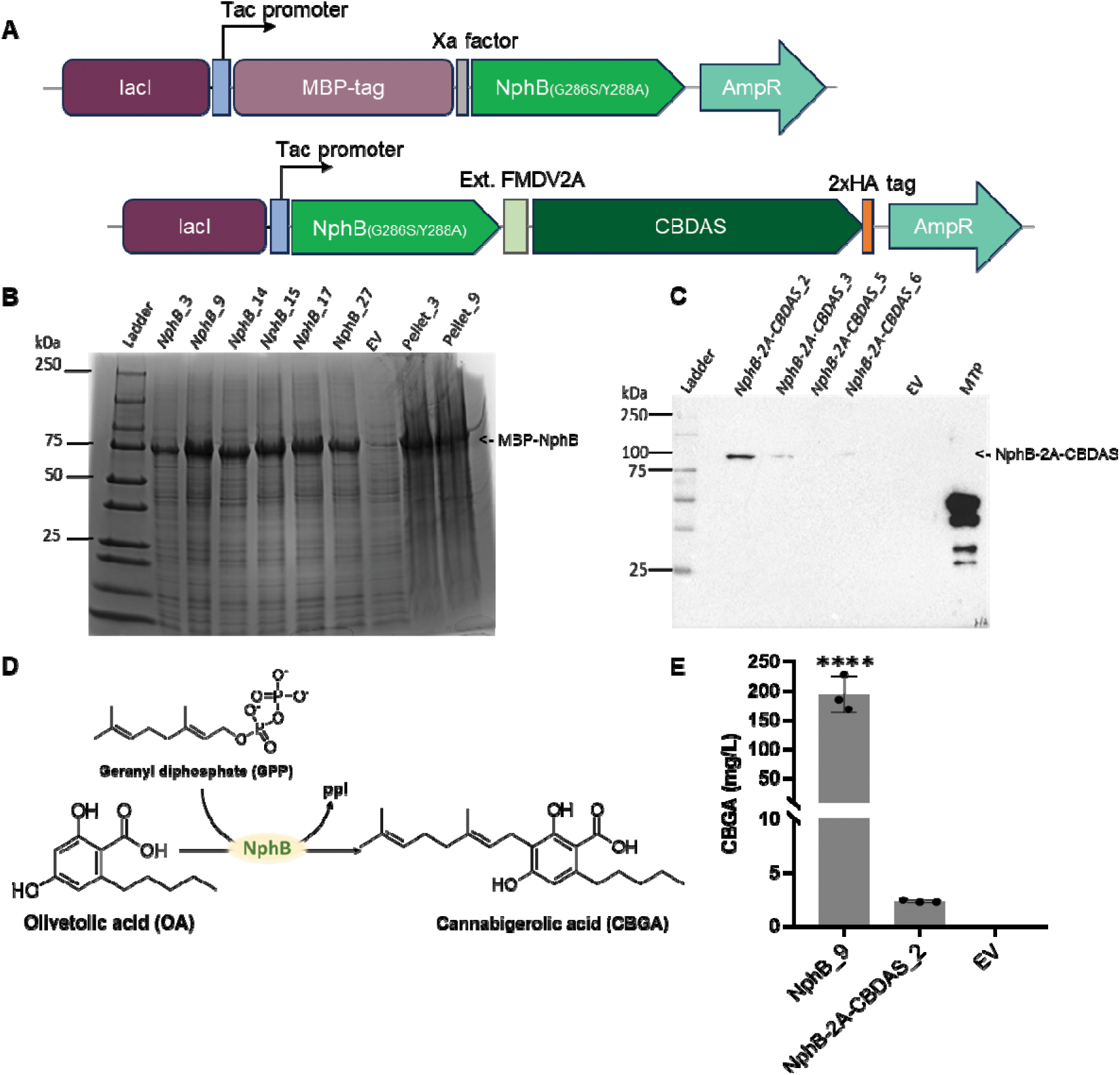
*NphB e*xpression and co-expression with *CBDAS* (*NphB-2A-CBDAS*) in *E. coli* led to the accumulation of the protein of both genes, but only NphB was functionally active. A) expression cassettes; B) Coomassie-stained SDS gel showing the NphB protein (fused to MBP-tag ≈ 75 kDa) accumulation compared to the EV; C) Western blot detection of *NphB-2A-CBDAS* (≈ 100 kDa) using HA antibody, the lower panel is the cropped Coomassie-stained SDS gel after transfer onto the PVDF membrane; D) NphB *in* vitro enzymatic reaction leading to the synthesis of CBGA from the C–C prenylation of olivetolic acid (OA) by geranyl diphosphate (GPP); E) CBGA concentration after *in* vitro enzymatic assays using NphB and NphB-2A-CBDAS protein extracts. CBGA quantification in different samples was determined using the standard curve plotted with the commercial CBGA standard used in the HPLC-DAD runs. EV: empty vector, MTP: Multi-Tag protein, 2A: FMDV2A. n=3, results are shown as mean ± standard deviation; p< 0.0001.

*E. coli* strain Rosetta BL21 (DE3) pLysS was transformed with the recombinant plasmids to express transgenes and produce proteins. Cell lysates were used to assess protein accumulation and enzymatic activity. When expressed alone, the NphB protein was detected by SDS-PAGE (**Figure 2B**). The fused protein NphB-2A-CBDAS was detected by western blot (WB) using an anti-HA antibody, appearing predominantly as the uncleaved fusion protein (**Figure 2C**).

CBGA and CBDA production was evaluated using GPP and OA as substrates (**Figure 2D)** and analyzed by HPLC-DAD, followed by confirmation with HPLC-MS/MS (**Figure S4 A-B**). The NphB enzyme showed *in vitro* activity in both constructs but produced significantly more CBGA when expressed alone (193 ± 31 mg/L) compared to co-expression with CBDAS (2.3 ± 0.1 mg/L) (**Figure 2E**, *p* < 0.0001). Although CBDAS protein accumulated, CBDA was not detected in the enzymatic assay, suggesting the enzyme was inactive (**Figure S4A**).

These results indicate that the codon-optimized *NphB* and *CBDAS* genes can be expressed and translated in *E. coli* under the tested conditions, with detectable levels of protein accumulation. However, cytosolic expression of *NphB* alone yields higher levels of a functional enzyme *in vitro*, whereas co-expression with *CBDAS* in a single ORF reduces protein accumulation and results in an inactive *CBDAS* enzyme.

### 3.2 Co-expression of *NphB* and *CBDAS* transgenes as a single ORF in the microalga *C. reinhardtii*

We hypothesized that using a eukaryotic photosynthetic heterologous host with demonstrated potential in terpene biosynthesis, such as *C. reinhardtii* [45, 55, 56], could facilitate CBDA production. As a first step, we used a construct encoding a bicistronic transgene *NphB-2A-CBDAS* (hereafter referred to as C1) driven by the *C. reinhardtii* fusion promoter ARp (a hybrid of the *Heat-Shock Protein 70A* and the *ribulose bisphosphate carboxylase/oxygenase small subunit* gene promoters) (**Figure 3**) [72].

**Figure 3.**
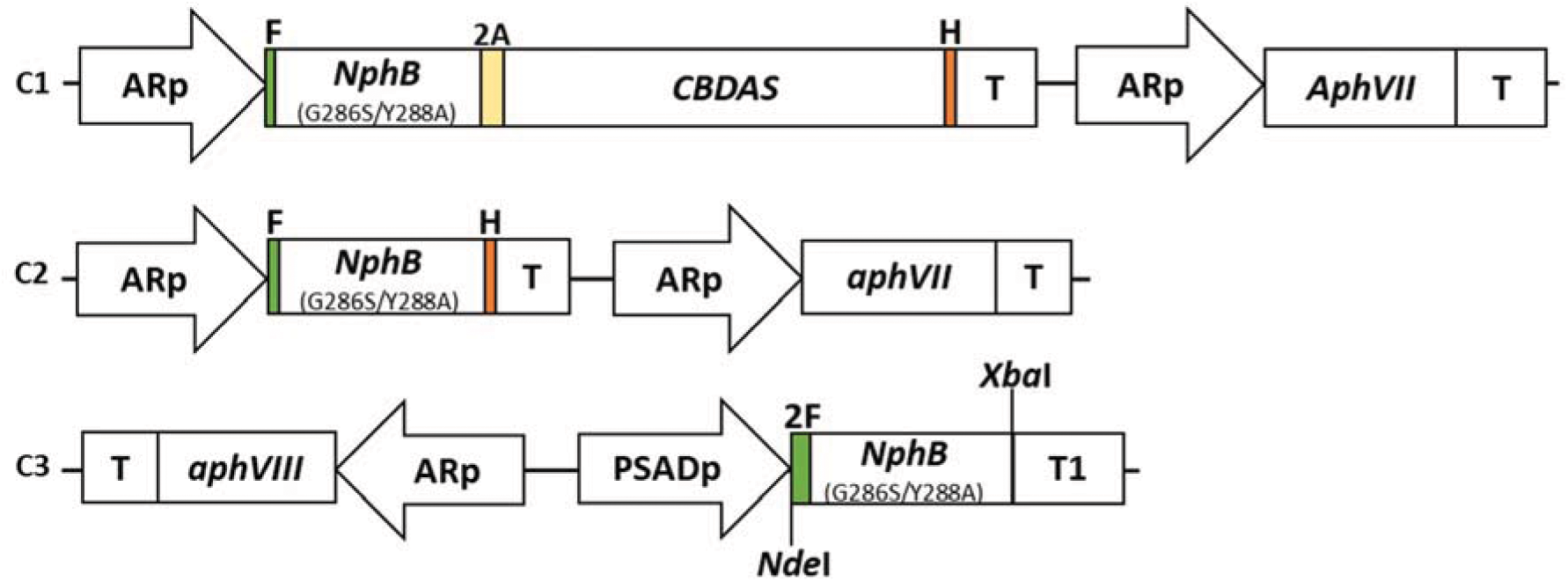
Genetic constructs used in the present study. **C1,** the bicistronic genetic construct containing the double-mutated soluble aromatic prenyl transferase gene (*NphB*_G286S/Y288A_) from *Streptomyces* sp. strain CL190 (Valliere et al. 2019) linked to the *C. sativa’s* cannabidiolic acid synthase gene (*CBDAS*) by the extended Foot and Mouth Disease Virus 2A (*extFMDV2A*) peptide sequence (Plucinak et al. 2015; Taura et al. 2007). **C2** is the monocistronic genetic construct harboring the *NphB* gene. Genes sequences of the C1 and C2 constructs were tagged in their N and C-terminal regions with Flag (F) and HA (H)-tags, respectively, and their expression was driven by the *C. reinhardtii* fusion promoter *HSP70A-RBCS2/5’UTR* (ARp) and its *3’UTR/RBCS2* terminator (T). The two constructs (C1 and C2) were cloned into the pOPt_*mRuby2* vector backbone harboring the Hygromycin B resistance gene (*aph*VII) as an antibiotic selection marker (Lauersen, Kruse, and Mussgnug 2015). **C3** is the monocistronic genetic construct with the *NphB* gene under the regulation of *C. reinhardtii* strong promoter *PSAD* (PSADp) and *PSAD* terminator (T1). In construct C3, the *NphB* sequence was 2xFlag (2F)-tagged in the N-terminal region, and the construct was cloned into the pSL18 vector backbone harboring the paromomycin resistance gene (*aph*VIII) under the regulation of ARp and T as an antibiotic selection marker (Fischer and Rochaix 2001).

Construct C1 was successfully integrated into the nuclear genome of the wild-type strain CC-125 by electroporation and into UVM4 and UVM11 by glass bead transformation. Transformation of CC-125 yielded approximately 3,000 colonies (**Figure S5A; Table 1**), while 100 and 198 colonies were obtained for UVM4 and UVM11, respectively (**Figure S13A and B; Table S6**). To evaluate transgene stability and integration, 384 randomly selected colonies from CC-125 transformants were subcultured under antibiotic selection for at least five rounds before further analysis. **Figure S5G** illustrates the high-throughput subculturing layout used for clone screening. PCR analysis of these 384 clones targeted the full-length expression cassette, spanning from the promoter to the terminator. Twenty-one clones (5.5%) yielded a PCR product of the expected size (3.5 kb), indicating successful integration (**Figure S6A; Table 1**). The integrity of the inserted cassette was further confirmed by Sanger sequencing. These PCR-positive (PCR) clones were cultivated in fresh TAP liquid medium with antibiotics and analyzed for transgene expression.

**Table 1:**
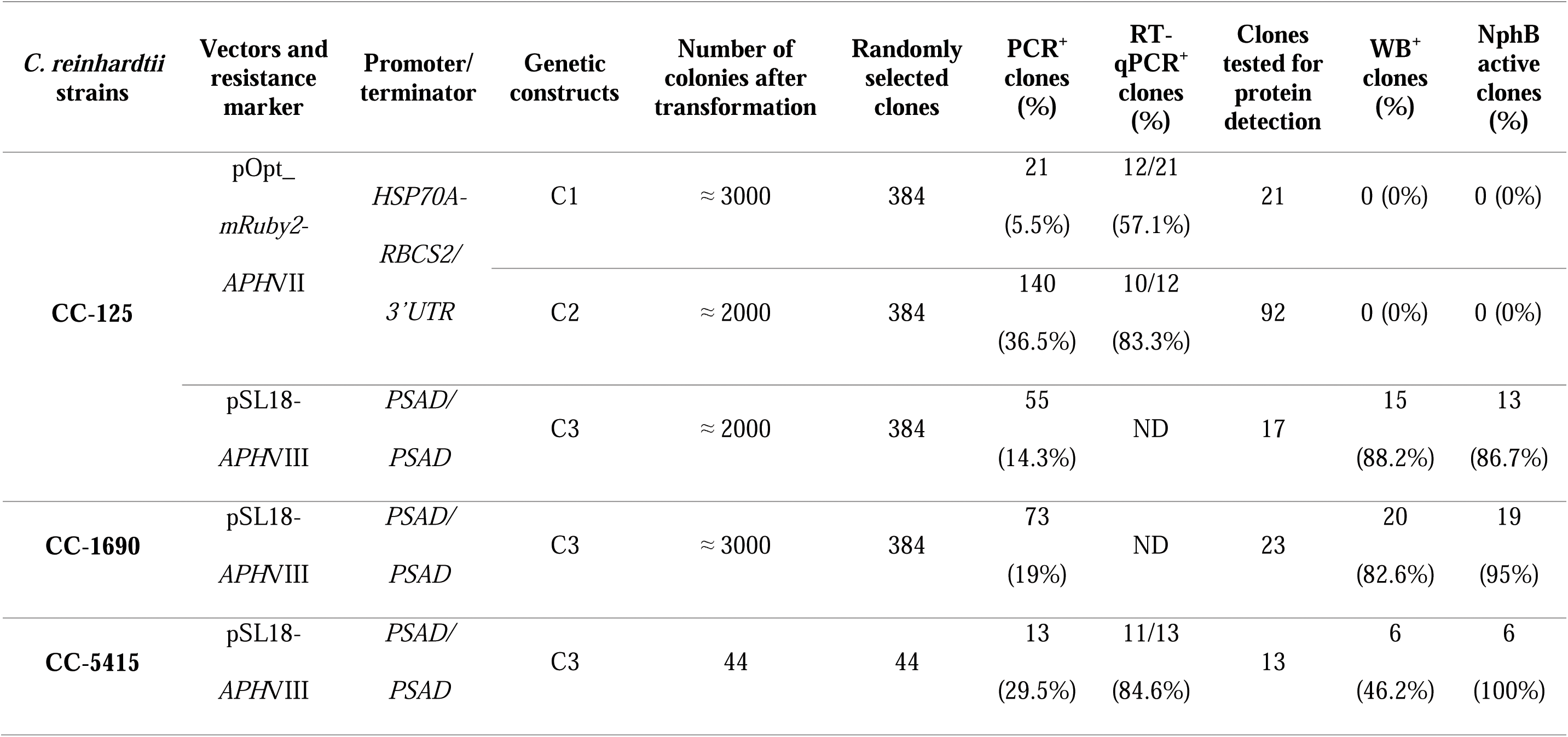
Summary of stable recombinant *C. reinhardtii* clones after five rounds of subculturing collated with the PCR-positive clones and the transformants exhibiting NphB activity *in vitro*, leading to the synthesis of CBGA.

| *C. reinhardtii* strains | Vectors and resistance marker | Promoter/terminator | Genetic constructs | Number of colonies after transformation | Randomly selected clones | PCR<sup>+</sup> clones (%) | RT-qPCR<sup>+</sup> clones (%) | Clones tested for protein detection | WB<sup>+</sup> clones (%) | NphB active clones (%) |
| --- | --- | --- | --- | --- | --- | --- | --- | --- | --- | --- |
| CC-125 | pOpt_<br>*mRuby2*-<br>*APHVII* | *HSP70A*-<br>*RBCS2*/<br>*3'UTR* | C1 | ≈ 3000 | 384 | 21<br>(5.5%) | 12/21<br>(57.1%) | 21 | 0 (0%) | 0 (0%) |
|  |  |  | C2 | ≈ 2000 | 384 | 140<br>(36.5%) | 10/12<br>(83.3%) | 92 | 0 (0%) | 0 (0%) |
|  | pSL18-<br>*APHVIII* | *PSAD*/<br>*PSAD* | C3 | ≈ 2000 | 384 | 55<br>(14.3%) | ND | 17 | 15<br>(88.2%) | 13<br>(86.7%) |
| CC-1690 | pSL18-<br>*APHVIII* | *PSAD*/<br>*PSAD* | C3 | ≈ 3000 | 384 | 73<br>(19%) | ND | 23 | 20<br>(82.6%) | 19<br>(95%) |
| CC-5415 | pSL18-<br>*APHVIII* | *PSAD*/<br>*PSAD* | C3 | 44 | 44 | 13<br>(29.5%) | 11/13<br>(84.6%) | 13 | 6<br>(46.2%) | 6<br>(100%) |

In *C. reinhardtii,* transgenic DNA is often subjected to epigenetic silencing mechanisms [50, 53, 70]. Therefore, stable genomic integration does not guarantee robust gene expression. To assess transcriptional activity, we measured the relative expression of *NphB-2A-CBDAS* mRNA using RT-qPCR, normalized to the *histone H3* housekeeping gene. Among the 21 PCR clones, 16 (76.2%) exhibited detectable transgene expression, whereas no expression was observed in the empty vector (EV) or wild-type (WT) controls (**Figure 4A**; **Table 1**). Notably, transgene expression levels varied substantially among clones, with transcript abundance in clone 21 being approximately 733-fold higher than in clone 16. This significant variability highlights the complexity of transgene regulation in *C. reinhardtii*, possibly due to transcriptional silencing or positional effects [48, 50, 53].

**Figure 4.**
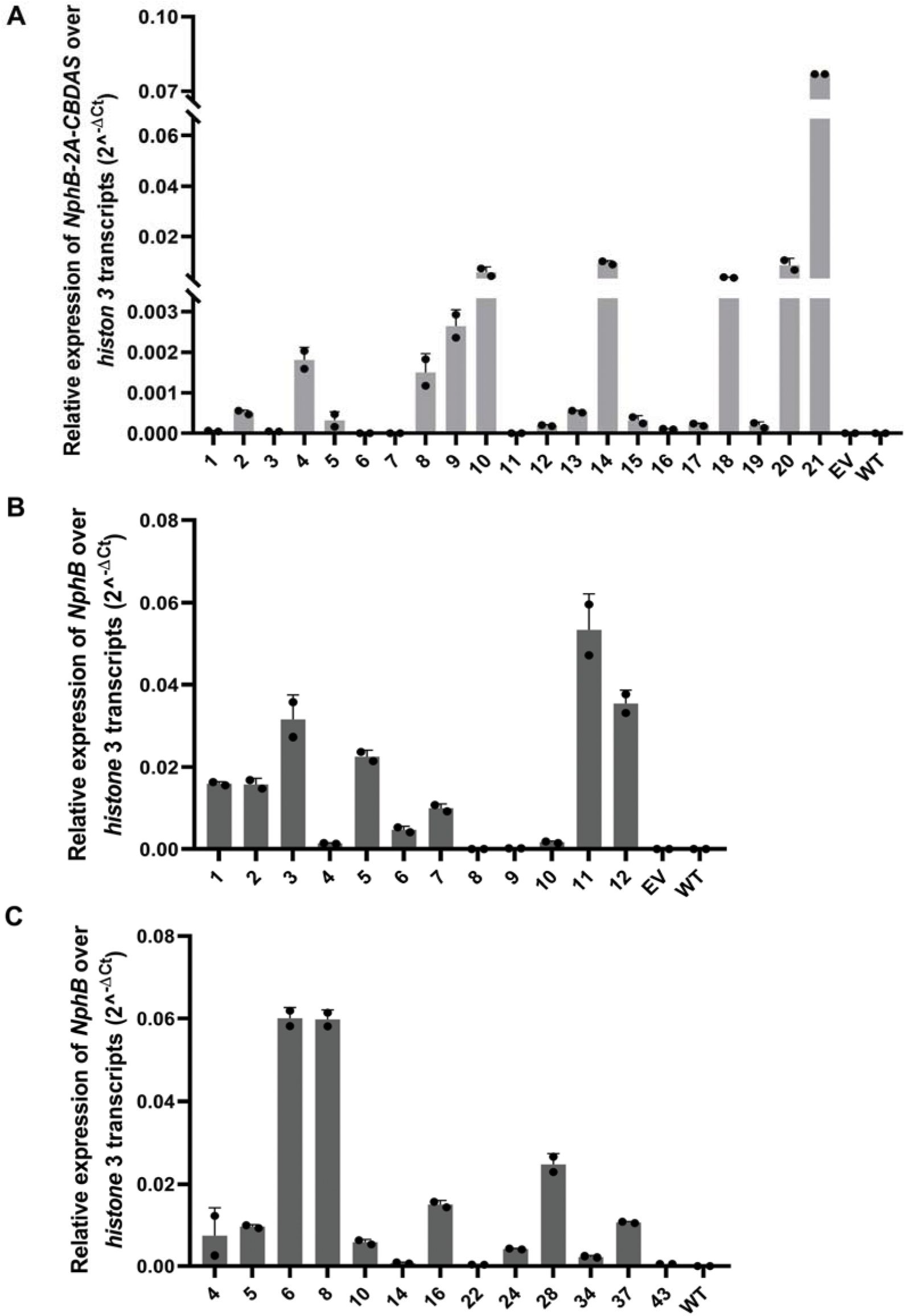
Relative expression of *NphB* and *CBDAS* over *histone* 3 transcripts. The clones tested from the PCR+ transformants generated with the genetic constructs C1, C2, and C3 show significant variation in the transgenes’ expression in clones from the same transformed cell lines. A) C1 expression in the strain CC-125, B) C2 expression in the strain CC-125, and C) C3 expression in the strain CC-5415. EV: empty vector; WT: wild-type. n=2, results are shown as mean ± standard deviation.

Despite these promising transcriptional results, protein accumulation was not detected by western blot (WB) in any of the tested transformants (n = 5), even though positive controls (purified multi-tagged proteins) were correctly identified (**Figure S7A**). No specific bands were observed at the expected sizes (∼37 kDa for NphB, ∼60 kDa for CBDAS, or ∼100 kDa for the fused protein). Consistently, *in vitro* enzymatic assays using total protein extracts from these clones with OA and GPP as substrates failed to produce CBGA or CBDA under the tested conditions. HPLC analysis of the enzymatic products showed no detectable cannabinoid peaks compared to the positive control (*NphB* expressed in *E. coli*) and the standard cannabinoid profiles (**Figure S8**).

We also evaluated C1 transformants of the high-expression strains UVM4 and UVM11. Although PCR analysis confirmed successful integration of the full-length transgene in both strains (**Figure S13C**), no protein accumulation was observed in any transformants by western blot (**Figure S13D and E**).

The absence of NphB and CBDAS protein accumulation in the *C. reinhardtii* transformants expressing the bicistronic construct C1 may be due to factors such as the large size of the transgenic DNA and potential regulatory complexity [48, 73, 74]. We, hypothesized that expressing the two genes separately, as monocistronic units, might help overcome these limitations.

### 3.3 Expression of *NphB* and *CBDAS* using independent ORFs

To test this, we split the original C1 construct into two independent constructs: C2 and C4. These constructs carried *NphB* and *CBDAS*, each tagged with *2xFlag* (N-terminal) or *2xHA* (C-terminal) epitopes. Both constructs were driven by the same *C. reinhardtii* AR promoter and terminator used in C1 (**Figure 3; Figure S7C**). Strain CC-125 was transformed with either C2 alone or a combination of C2 and C4 to evaluate single and co-expression scenarios.

Following transformation with C2, approximately 2,000 colonies were obtained on selective plates (**Figure S5B; Table 1**). A subset of 384 colonies was randomly selected and subcultured through five passages on TAP medium supplemented with 15 mg/L hygromycin. PCR screening identified 140 positive clones (36.5%) harboring the complete expression cassette (**Figure S6B; Table 1**).

RT-qPCR analysis of selected C2 transformants revealed variable levels of *NphB* mRNA expression among clones from the same transformation line (**Figure 4B**). Specifically, 10 out of 12 tested clones (83.3%) showed detectable *NphB* transcript levels, while expression was absent in wild-type (WT) and empty vector (EV) controls (**Figure 4B**; **Table 1**). When co-transforming C2 and C4, *NphB* and *CBDAS* expression patterns were comparable to those observed in C1 and C2 alone (**Figure S7C**).

Despite the transcriptional activity, western blot analysis revealed no detectable accumulation of the expected proteins. As with the C1 construct, only non-specific bands were observed using the anti-Flag antibody, and no bands corresponding to the expected molecular weights of NphB or CBDAS were detected (**Figure S7B and D**). Additionally, *in vitro* enzymatic assays performed with protein extracts from the transformants failed to yield CB metabolites, corroborating the lack of functional protein accumulation.

These results indicate that the NphB and CBDAS coding sequences were successfully integrated into the nuclear genome and transcribed, but did not lead to detectable levels of functionally active enzymes in our assays. This suggests that post-transcriptional silencing or translational repression may be affecting protein expression. Known variables such as the host strain, promoter and terminator sequences, or transgene sequence context in *C. reinhardtii* may contribute to these effects [48, 75].

### 3.4 PSAD-driven nuclear expression enables functional accumulation of NphB but not CBDAS in *C. reinhardtii*

Given the repeated absence of detectable CBDAS protein and enzymatic activity under various genetic configurations, including constructs driven by the *PSAD* promoter and terminator, we excluded CBDAS from further optimization efforts and focused exclusively on enhancing NphB expression. These adverse outcomes strongly suggest that CBDAS expression in *C. reinhardtii* under the tested conditions remains a significant bottleneck and warrants independent investigation beyond the scope of this study.

We therefore constructed a new expression cassette (C3) in which the codon-optimized *NphB* gene was placed under the control of the *C. reinhardtii* nuclear *PSAD* promoter and terminator (**Figure 3**). This cassette was introduced by electroporation into the nuclear genome of the wild-type CC-125 strain. Six to seven days post-transformation, approximately 2,000 colonies were obtained on TAP agar plates supplemented with 15 mg/L paromomycin (**Figure S5D; Table 1**). As with previous constructs, 384 colonies were randomly selected and subcultured through five rounds of antibiotic selection. Colony PCR screening identified 55 transformants (14.3%) with a fully integrated expression cassette (**Table 1**).

To evaluate transgene expression and protein accumulation, 23 of the PCR+ clones were cultured in 25 mL TAP medium for 5 days (late exponential growth phase). Among these, 17 maintained stable transgene integration (**Figure S6C; Table 1**). Western blot analysis revealed that 15 out of 17 clones (88.2%) produced a protein band of ∼37 kDa, consistent with the expected size of NphB (**Figure 5A**; **Table 1**). These results confirm that the *PSAD* promoter and terminator efficiently drive transgene expression in *C. reinhardtii*, under the present experimental conditions, enabling accumulation of detectable, functional NphB.

**Figure 5.**
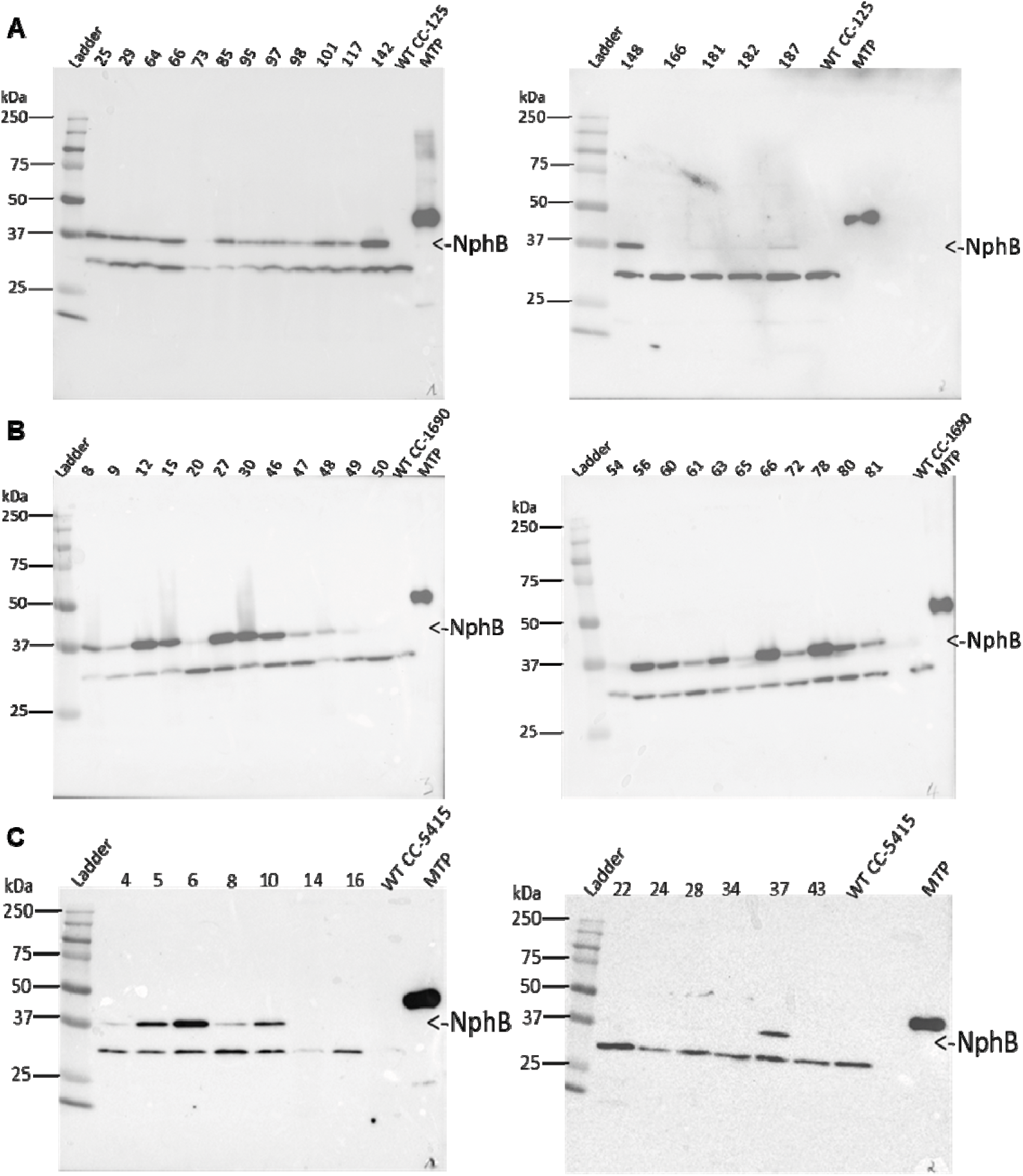
Expressing NphB under the PSAD regulation led to the accumulation of the protein of interest. In different WT strains of C*. reinhardtii*, Western blot (WB) detected the protein of interest on the membrane after incubation with the Flag antibody. A) NphB detection in 13 clones among the 15 PCR+ clones tested from the strain CC-125; B) NphB detection in 20 clones among the 23 PCR+ clones tested from the strain CC-1690; C) NphB detection in 6 clones among the 13 PCR+ clones tested from the strain CC-5415. *NphB’s* expected size is ≈ 37 kDa; MTP: multi-tag protein used as a positive control for the antibody treatment; WT: wild type.

In addition to the wild-type strain CC-125, we selected CC-1690 and CC-5415 to assess the reproducibility of results across different wild-type genetic backgrounds. This approach allowed us to determine whether promoter and construct performance observed in CC-125 would be consistent in other genetic contexts.

### 3.5 *NphB* expression cassette integration in different *C. reinhardtii* strains

To investigate whether the genetic background of *C. reinhardtii* influences the expression of heterologous genes, we introduced the C3 expression cassette, harboring *NphB* under the control of the *PSAD* promoter and terminator, into two additional wild-type strains: CC-1690 and CC-5415. After electroporation, we observed notable differences in transformation efficiency and protein expression across strains. Indeed, approximately 3,000 colonies were obtained for CC-1690 and only 44 for CC-5415, compared to about 2,000 for CC-125 (**Figure S5D-F; Table 1**). PCR screening of transformants revealed successful integration of the full-length C3 cassette in 19.0% (73/384) of CC-1690 clones and 29.5% (13/44) of CC-5415 clones, compared to 14.3% (55/384) in CC-125 (**Figure S6C-E; Table 1)**.

Quantification of *NphB* mRNA expression relative to the housekeeping gene *histone 3* in CC-5415 PCR+ clones showed expression profiles consistent with those previously observed for C2 (**Figure 4C**, **Table 1**). Western blot analysis confirmed the accumulation of the 2xFlag-NphB protein (∼37 kDa) in 82.6% (20/23) of CC-1690 and 46.2% (6/13) of CC-5415 transformants. This contrasts with the 88.2% (15/17) of CC-125 clones exhibiting detectable NphB protein (**Figure 5 A-C**; **Table 1**).

These results demonstrate that NphB can be functionally expressed in all three *C. reinhardtii* strains tested, although expression efficiency and protein accumulation levels vary. This strain-dependent variability highlights the importance of host background, such as high-expression strain UVM4, when optimizing nuclear transgene expression in *C. reinhardtii*.

### 3.6 *C. reinhardtii*-produced NphB Catalyzes CBGA Production *in vitro*

To investigate the ability of *C. reinhardtii*-expressed NphB to catalyze CBGA production, we first attempted *in vivo* synthesis by supplementing the culture medium with 2 mM olivetolic acid (OA) at the late exponential growth phase. Metabolites were extracted 24 hours later and analyzed by HPLC-MS/MS (**Figures S12A and B**). Despite confirmed NphB protein accumulation in transformants, CBGA was not detected under these conditions.

We therefore conducted *in vitro* enzymatic assays using the soluble protein fraction from transformants, with OA and geranyl diphosphate (GPP) as substrates. Given the number of WB^+^ transformants (n = 41) and the cost of substrates, we implemented an initial screening using a single replicate per clone and 0.5 mM of each substrate to identify candidates exhibiting NphB activity. CBGA production was detected in 38 clones, as evidenced by a signal at a retention time of 8.309 min consistent with the CBGA standard and a positive control containing recombinant NphB expressed in *E. coli* (**Figures S8 and S9**). This compound was identified as CBGA based on its molecular ion (m/z 361) and fragmentation pattern, confirmed by HPLC-MS/MS (**Figure S10**). No CBGA signal was observed in WT protein reactions.

Among the strains tested, CC-1690 and CC-5415 showed slightly higher proportions of active clones, with 19/20 (95%) and 6/6 (100%) clones, respectively, exhibiting NphB activity, compared to 13/15 (86.7%) for CC-125 (**Table 1, Table S5**). The clones with the highest NphB activity from each strain, clones 12 (CC-1690), 25 (CC-125), and 37 (CC-5415), were selected for confirmation in triplicate. Western blot confirmed NphB accumulation in all three (**Figure 6A**). In contrast, no NphB signal was detected in extracts from negative controls (EV, WT) or clones 21 and 5 from constructs C1 and C2, respectively, consistent with previous findings (**sections 3.2** and **3.3**).

**Figure 6.**
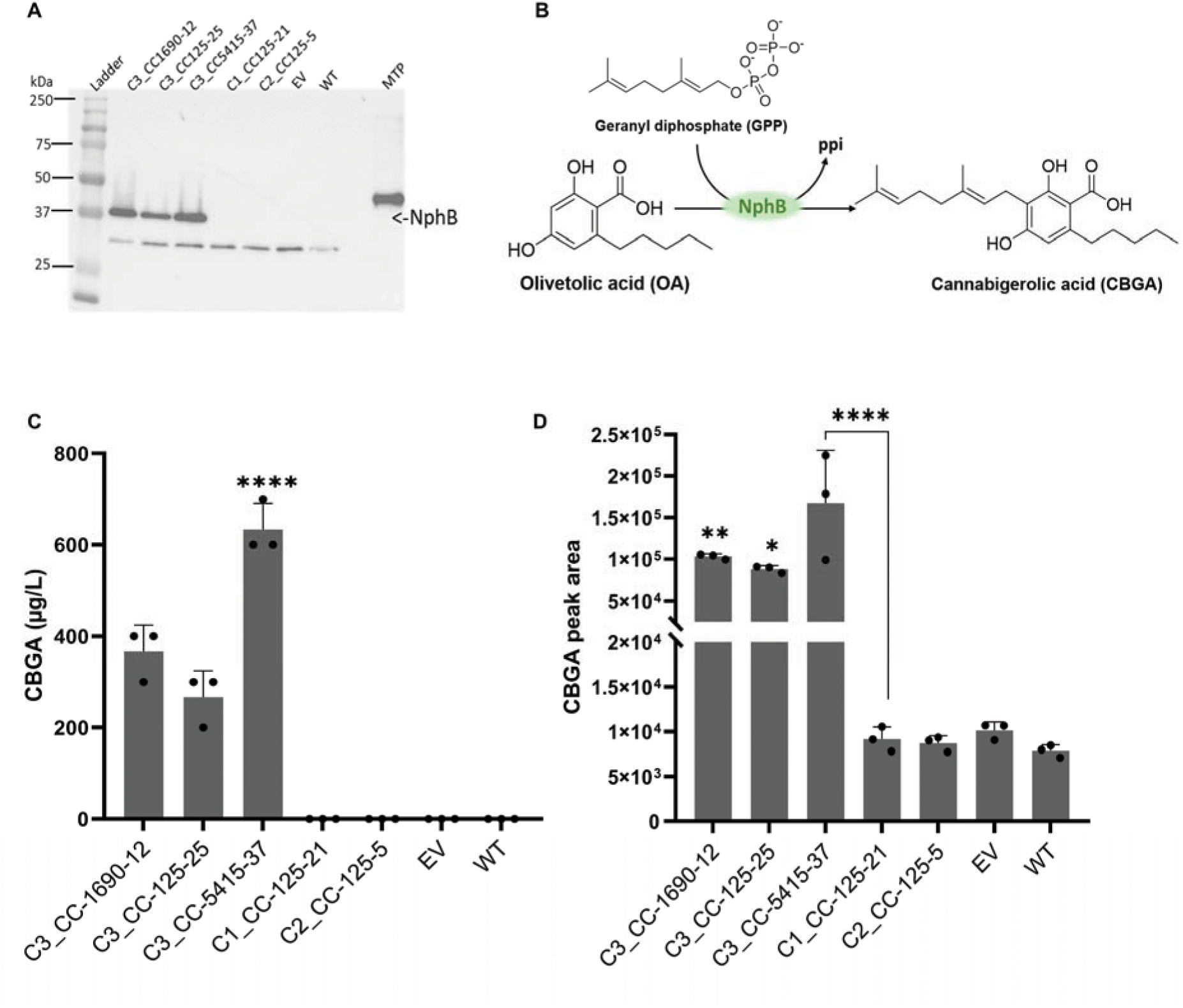
*C. reinhardtii* successfully expressed the *NphB* transgene under the PSAD regulation, accumulating active proteins. A) Western blot showing NphB detection in clones 12, 25, and 37, respectively, selected from strains CC-1690, 125, and 5415, compared to the negative control EV and WT, but also to clones 21 and 5, respectively selected from C1 and C2 transformants expressing NphB-2A-CBDAS and NphB under AR promoter and terminator regulation; B) NphB *in* vitro enzymatic reaction leading to the synthesis of CBGA from the C–C prenylation of olivetolic acid (OA) by geranyl diphosphate (GPP); C) CBGA concentration obtained after *in* vitro enzymatic assays using proteins crude extracts from the *C. reinhardtii* transformants expressing *NphB*, analyzed and quantified by HPLC-DAD using commercial CBGA standard curve; D) HPLC-MS/MS confirmation of CBGA production generating peak areas proportional to CBGA concentration in each sample. EV: empty vector, MTP: Multi-Tag protein, 2A: FMDV2A, WT: wild type. n=3, results are shown as mean ± standard deviation.

HPLC-DAD analysis further validated NphB enzymatic activity by detecting a product with a retention time of 10.146 min, matching the CBGA standard (**Figure S11**). Increasing substrate concentrations (OA and GPP) significantly enhanced CBGA yields, resulting in 2.6- to 6.3-fold higher production compared to assays using 0.5 mM substrates, which were below the HPLC-DAD detection threshold of 100 µg/L (**Figure 6B**; **Table S5**). Clone 37 produced the highest CBGA titer (633 ± 58 µg/L), followed by clones 12 (367 ± 58 µg/L) and 25 (267 ± 58 µg/L) (**Figure 6C**). In contrast, no CBGA signal above background was detected in extracts from clones 5 and 21, or the EV and WT controls.

Statistical analysis using one-way ANOVA, followed by Tukey’s post-hoc test, confirmed that CBGA peak areas for the WB^+^ clones (12, 25, and 37) were significantly higher (p < 0.0001) than those from WB^−^ clones and negative controls (**Figure 6D**). Altogether, these findings provide, to our knowledge, the first reported evidence of functionally active NphB enzyme production in the nuclear genome of *C. reinhardtii*, which can catalyze CBGA synthesis *in vitro*. Despite the absence of detectable CBGA accumulation *in vivo*, the successful *in vitro* enzymatic activity supports the biological functionality of the heterologously expressed NphB. These results provide a foundation for future optimization strategies to enhance precursor availability, refine subcellular targeting, and improve metabolic compartmentalization, key factors that will be discussed in the following section.

## 4. Discussion

This study provides initial evidence that *C. reinhardtii* can serve as a host for heterologous expression of genes involved in cannabinoid biosynthesis. In particular, we achieved functional expression of the prenyltransferase *NphB* in the cytosol, with protein accumulation dependent on the promoter/terminator pair. However, consistent with prior studies, *CBDAS* expression in the cytosol failed to yield active protein [21], highlighting a significant bottleneck in reconstituting the complete cannabinoid pathway in *C. reinhardtii*.

Our bicistronic and monocistronic constructs, encoding a codon-optimized NphB (G286S/Y288A mutant) and CBDAS from *Cannabis sativa* [43, 63], were first validated in *E. coli*, where NphB showed robust CBGA production (up to 193 ± 31 mg/L) in the monocistronic configuration (**Figure 2E**). Conversely, the co-expression with CBDAS (bicistronic construct) led to lower CBGA titers (2.3 ± 0.1 mg/L), possibly due to incomplete 2A peptide cleavage or reduced *NphB* expression [16, 76, 77]. Notably, while CBDAS protein was detectable as a fused protein to NphB, no CBDA was synthesized *in vitro*, supporting previous observations that CBDAS functionality may depend on eukaryotic expression systems with proper folding, glycosylation, and subcellular localization [21, 31, 78–80].

To express these genes in *C. reinhardtii*, we introduced various constructs into the nuclear genome of multiple wild-type strains (CC-125, CC-1690, CC-5415), including the high-expression strains UVM4 and UVM11. Under the HSP70A-RBCS2 (AR) promoter, transgene transcription occurred but did not result in detectable NphB or CBDAS proteins, indicating post-transcriptional silencing or insufficient mRNA processing [48, 81]. Switching to the PSAD promoter and terminator resulted in 88.2% of transformants (in CC-125) accumulating NphB protein in the cytosol, enabling CBGA biosynthesis *in vitro* up to 633 ± 58 µg/L under our assay conditions (**Figure 6C**). This pattern was reproducible in the other wild-type strains tested, supporting the robustness of PSAD regulatory elements for expressing this transgene [61, 75, 82].

Nevertheless, the *in vivo* production of CBGA in *C. reinhardtii* remains elusive. Even upon exogenous olivetolic acid supplementation, no CBGA was detected in culture media or biomass. This suggests potential limitations in precursor availability, notably the cytosolic pool of geranyl pyrophosphate (GPP) or inefficient uptake and conversion of olivetolic acid. Similar issues have been reported in other algal systems such as *Phaeodactylum tricornutum* engineered for cannabinoid production [22, 23]. Strategies to overcome this limitation include overexpression of genes in the MEP pathway (e.g., *DXS*, *DXR*, *HDR*), suppression of native GPP-consuming pathways (e.g., competing terpene synthases), or employing synthetic scaffolds to enhance substrate channeling [17, 55, 83–85]. In addition, varying culture conditions during olivetolic acid supplementation, including light intensity, nutrient levels, and timing, may improve precursor assimilation and enzymatic conversion, offering a practical route to *in vivo* CBGA production [22, 86].

On the other hand, the persistent failure to detect CBDAS protein or CBDA further highlights the need for subcellular targeting strategies. ER or vacuolar localization, or secretion, may enhance CBDAS folding and function, as demonstrated in *Phaeodactylum tricornutum*, yeast, and *Nicotiana benthamiana* [21, 31, 78–80, 87], or with other recombinant proteins in *C. reinhardtii* [88, 89]. Fusion with ER signal peptides or retention sequences (e.g., HDEL) should be tested [90–92]. Beyond the ER, the vacuole may also serve as a promising destination for CBDAS, offering a compartment with a distinct redox environment and reduced protease activity that could improve enzyme stability and preserve catalytic function [14, 17, 78, 93]. Furthermore, since cannabinoid synthases like CBDAS and THCAS require flavin adenine dinucleotide (FAD) as a cofactor for enzymatic activity [3, 94], ensuring adequate FAD availability or cofactor-binding efficiency in the heterologous host may help restore functional activity and should be considered in future engineering strategies.

One crucial factor that may underlie inconsistent or failed expression of nuclear transgenes in *C. reinhardtii* is incorrect mRNA processing [53]. Nuclear-expressed transgenes often suffer from inefficient splicing, poor transcript stability, or defective nuclear export, especially when using intronless coding sequences [48, 51, 95, 96], as was the case for both NphB and CBDAS in our study. The absence of introns can result in mRNA not being adequately recognized by the splicing machinery, leading to nuclear retention or degradation [51, 52, 97]. In particular, CBDAS transcripts may have been transcribed but failed to be exported or translated due to such processing errors. Cryptic splice sites introduced during codon optimization or weak 3′ UTR sequences may have contributed to aberrant mRNA forms or destabilization [47, 48, 75, 96]. These possibilities highlight the potential importance of intron incorporation [51, 96, 98], validated terminators [48, 75], and transcript-stabilizing elements in future construct designs to ensure robust mRNA maturation and expression.

Promoter choice also emerged as a critical factor [99–101]. While the AR promoter was sufficient for mRNA production, it failed to yield detectable protein in our constructs (**Figure S7 A-D**), even when using the high-expression strains UVM4 and UVM11 for the expression of C1 in our experimental conditions (**Figure S13 D and E**). In contrast, the PSAD promoter and terminator proved highly effective for NphB expression (**Figure 5A-C**). Future designs should test synthetic or hybrid promoters and alternative terminators with more potent transcript stabilization properties [100, 102]. Chromatin context and transgene insertion site variability must also be addressed via site-specific integration tools or insulators [47, 48, 50, 70, 103].

Our results also indicate that multiple *C. reinhardtii* wild-type strains (CC-125, CC-1690, CC-5415) can support heterologous expression of prenyltransferases under the tested conditions. This expands the toolbox for strain selection and suggests that future efforts may benefit from strain-specific optimizations targeting cell wall properties, transformation method and efficiency, and endogenous protease activity [51–53, 70, 93, 104]. Although NphB and CBDAS have been previously expressed in phtosynthetic marine diatoms, such as *P. tricornutum* [21, 22], this study is the first to report the successful nuclear expression and functional activity of the bacterial aromatic prenyltransferase NphB in the green microalga *Chlamydomonas reinhardtii*, enabling *in vitro* CBGA biosynthesis. This represents a key advancement given the photosynthetic nature, plant-like protein processing machinery, and regulatory complexities of *C. reinhardtii*, which differ substantially from those of marine diatoms, highlighting both the advantages of this green alga as a plant-like host and the importance of studying multiple microalgal platforms until robust cannabinoid production is achieved [44, 47].

While this work represents a first step towards cannabinoid biosynthesis in *C. reinhardtii*, several significant limitations must be acknowledged. CBGA production was only demonstrated *in vitro*; no detectable *in vivo* accumulation was achieved despite supplementation with olivetolic acid, indicating potential limitations in precursor availability, uptake, or intracellular compartmentalization. CBDAS transcripts were detected, but no functional protein was observed, suggesting challenges with translation efficiency, post-translational modifications, or subcellular targeting. Our constructs used intronless coding sequences, which in *C. reinhardtii* may result in suboptimal mRNA processing and stability. Protein yields and enzyme activity varied between strains and constructs, highlighting the influence of host genetic background and promoter choice. Addressing the present challenges, through precursor supply, subcellular targeting, intron-dependent expression, and cofactor balance, will be critical to unlock its potential. Beyond its findings, the strategies tested in the present manuscript can extend to other metabolites, such as terpenoids. This work contributes to the broader field of synthetic biology and metabolic engineering by illustrating both the potential and the current limitations of the green microalga as a production host. While yeast, *E. coli*, and *Nicotiana benthamiana* have achieved heterologous cannabinoid production [2, 37, 79, 87], *C. reinhardtii* offers distinct advantages as a light-driven, carbon-fixing system with sustainability benefits [14, 45, 83, 99]. Emerging synthetic biology tools, including modular pathway assembly, synthetic scaffolds for substrate channeling, and CRISPR-based site-specific integration, provide promising strategies to improve yield and stability [83, 103, 105, 106]. At the bioprocess level, scalable photobioreactor systems could couple cannabinoid biosynthesis to renewable light and CO inputs, paving the way for environmentally sustainable production [83].

## Credit authorship contribution statement

**Serge Basile Nouemssi:** Conceptualization, Data curation, Formal analysis, Investigation, Methodology, Validation, Writing-original draft, Writing-review & editing. **Ayoub Bouhadada:** Conceptualization, Formal analysis, Investigation, Methodology, Writing-review & editing. **Remy Beauchemin:** Conceptualization, Methodology, Writing-review & editing. **Alexandre Custeau:** Formal analysis, Validation, Writing-review & editing. **Sarah-Eve Gelinas:** Formal analysis, validation, Writing-review & editing. **Natacha Merindol:** Conceptualization, Methodology, investigation, Data curation, Project administration, Writing-review & editing. **Fatma Meddeb-Mouelhi:** Conceptualization, Methodology, Project administration, Resources, Writing-review & editing. **Hugo Germain:** Project administration, Resources, Supervision, Writing-review & editing. **Isabel Desgagne-Penix:** Conceptualization, Visualization, Funding acquisition, Project administration, Resources, Supervision, Writing-review & editing.

## Acknowledgments

The authors wish to thank Dr. Mather Carscallen and the lab members past and present, specifically Drs Manel Ghribi and Bharat Bhusan Majhi, and Melodie B. Plourde for samples, technical support and helpful discussions.

## Supplementary data

Additional files and supplementary data to this article can be found at …

## References

1. Andre, C.M., J.-F. Hausman, and G. Guerriero, Cannabis sativa: the plant of the thousand and one molecules. Frontiers in Plant Science, 2016. 7: p. 174167.

2. Gu lck, T., et al., Synthetic Biology of Cannabinoids and Cannabinoid Glucosides in Nicotiana benthamiana and Saccharomyces cerevisiae. Journal of Natural Products, 2020. 83(10): p. 2877–2893.

3. Romero, P., et al., Comprehending and improving cannabis specialized metabolism in the systems biology era. Plant Science, 2020. 298: p. 110571.

4. Abd-Nikfarjam, B., et al., Cannabinoids in neuroinflammatory disorders: Focusing on multiple sclerosis, Parkinsons, and Alzheimers diseases. Biofactors, 2023. 49(3): p. 560–583.

5. Al-Ghezi, Z.Z., et al., *Combination of cannabinoids*, Δ*9-tetrahydrocannabinol and cannabidiol, ameliorates experimental multiple sclerosis by suppressing neuroinflammation through regulation of miRNA-mediated signaling pathways*. Frontiers in immunology, 2019. 10: p. 466430.

6. van den Hoogen, N.J., et al., Cannabinoids in chronic pain: therapeutic potential through microglia modulation. Frontiers in Neural Circuits, 2022. 15: p. 816747.

7. Abrams, D.I., The therapeutic effects of Cannabis and cannabinoids: An update from the National Academies of Sciences, Engineering and Medicine report. European journal of internal medicine, 2018. 49: p. 7–11.

8. Hazekamp, A., et al., The medicinal use of cannabis and cannabinoids—an international cross-sectional survey on administration forms. Journal of psychoactive drugs, 2013. 45(3): p. 199–210.

9. Dorbian. *Global Legal Cannabis Market Expected To Reach $58 Billion In* 2028. 2024 30 May 2024]; Available from: https://www.forbes.com/sites/irisdorbian/2024/03/07/global-legal-cannabis-market-expected-to-reach-58-billion-in-2028/?sh=465318e11d84.

10. Gülck, T. and B.L. Møller, Phytocannabinoids: origins and biosynthesis. Trends in plant science, 2020. 25(10): p. 985–1004.

11. Atakan, Z., Cannabis, a complex plant: different compounds and different effects on individuals. Therapeutic advances in psychopharmacology, 2012. 2(6): p. 241–254.

12. Shultz, Z.P., et al., Enantioselective Total Synthesis of Cannabinoids A Route for Analogue Development. Organic letters, 2018. 20(2): p. 381–384.

13. Dattatraya H. Dethe, R.D.E., Samarpita Mahapatra, Saikat Das and Vijay Kumar B., Protecting group free enantiospecific total syntheses of structurally diverse natural products of the tetrahydrocannabinoid family. Chemical communications, 2015. 51(14): p. 2871–2873.

14. Bolaños-Martínez, O.C., et al., Harnessing the advances of genetic engineering in microalgae for the production of cannabinoids. Critical Reviews in Biotechnology, 2022: p. 1–12.

15. Awwad, F., et al., Bioengineering of the Marine Diatom Phaeodactylum tricornutum with Cannabis Genes Enables the Production of the Cannabinoid Precursor, Olivetolic Acid. International Journal of Molecular Sciences, 2023. 24(23): p. 16624.

16. Zirpel, B., et al., Engineering yeasts as platform organisms for cannabinoid biosynthesis. Journal of biotechnology, 2017. 259: p. 204–212.

17. Luo, X., et al., Complete biosynthesis of cannabinoids and their unnatural analogues in yeast. Nature, 2019. 567(7746): p. 123–126.

18. Tan, Z., J.M. Clomburg, and R. Gonzalez, Synthetic pathway for the production of olivetolic acid in Escherichia coli. ACS Synthetic Biology, 2018. 7(8): p. 1886–1896.

19. Taura, F., et al., *Production of* Δ*1-tetrahydrocannabinolic acid by the biosynthetic enzyme secreted from transgenic Pichia pastoris*. Biochemical and biophysical research communications, 2007. 361(3): p. 675–680.

20. Zhang, Y., et al., Development of an efficient yeast platform for cannabigerolic acid biosynthesis. Metabolic Engineering, 2023. 80: p. 232–240.

21. Fantino, E., et al., Extrachromosomal expression of functional Cannabis sativa cannabidiolic acid synthase in Phaedodactylum tricornutum. Algal Research, 2025. 85: p. 103889.

22. Fantino, E., et al., Bioengineering Phaeodactylum tricornutum, a marine diatom, for cannabinoid biosynthesis. Algal Research, 2024: p. 103379.

23. Sene, N., et al., Impact of heterologous expression of Cannabis sativa tetraketide synthase on Phaeodactylum tricornutum metabolic profile. Biotechnology for Biofuels and Bioproducts, 2025. 18(1): p. 42.

24. Gagne, S.J., et al., Identification of olivetolic acid cyclase from Cannabis sativa reveals a unique catalytic route to plant polyketides. Proceedings of the National Academy of Sciences, 2012. 109(31): p. 12811–12816.

25. Sirikantaramas, S., et al., *The gene controlling marijuana psychoactivity: molecular cloning and heterologous expression of* Δ*1-tetrahydrocannabinolic acid synthase from Cannabis sativa L*. Journal of Biological Chemistry, 2004. 279(38): p. 39767–39774.

26. Sirikantaramas, S., et al., Tetrahydrocannabinolic Acid Synthase, the Enzyme Controlling Marijuana Psychoactivity, is Secreted into the Storage Cavity of the Glandular Trichomes. Plant and Cell Physiology, 2005. 46(9): p. 1578–1582.

27. Taura, F., S. Morimoto, and Y. Shoyama, Purification and characterization of cannabidiolic-acid synthase from Cannabis sativa L.: Biochemical analysis of a novel enzyme that catalyzes the oxidocyclization of cannabigerolic acid to cannabidiolic acid. Journal of Biological Chemistry, 1996. 271(29): p. 17411–17416.

28. Taura, F., et al., Phytocannabinoids in Cannabis sativa: recent studies on biosynthetic enzymes. Chemistry & biodiversity, 2007. 4(8): p. 1649–1663.

29. ElSohly, M. and W. Gul, Constituents of Cannabis sativa. Handbook of cannabis, 2014. 3(1093): p. 187–188.

30. Hanuš, L.O., et al., Phytocannabinoids: a unified critical inventory. Natural product reports, 2016. 33(12): p. 1357–1392.

31. Schmidt, C., M. Aras, and O. Kayser, Engineering cannabinoid production in Saccharomyces cerevisiae. Biotechnology Journal, 2024. 19(2): p. 2300507.

32. Hussain, T., et al., Cannabis sativa research trends, challenges, and new-age perspectives. Iscience, 2021. 24(12).

33. Kearsey, L.J., et al., Structure of the Cannabis sativa olivetol producing enzyme reveals cyclization plasticity in type III polyketide synthases. The FEBS Journal, 2020. 287(8): p. 1511–1524.

34. Degenhardt, F., F. Stehle, and O. Kayser, The biosynthesis of cannabinoids, in Handbook of cannabis and related pathologies. 2017, Elsevier. p. 13–23.

35. Carvalho, Â., et al., Designing microorganisms for heterologous biosynthesis of cannabinoids. FEMS yeast research, 2017. 17(4).

36. Krogh, A., et al., Predicting transmembrane protein topology with a hidden Markov model: application to complete genomes. Journal of molecular biology, 2001. 305(3): p. 567–580.

37. Ayakar, S.R., et al., Metabolic Engineering of E. coli for the Biosynthesis of Cannabinoid Products. 2020, Google Patents.

38. Page, J. and Z. Boubakir, Aromatic prenyltransferase from Cannabis. 2012. US 20120144523. 2018.

39. Bonitz, T., et al., Evolutionary relationships of microbial aromatic prenyltransferases. PloS one, 2011. 6(11): p. e27336.

40. Kuzuyama, T., J.P. Noel, and S.B. Richard, Structural basis for the promiscuous biosynthetic prenylation of aromatic natural products. Nature, 2005. 435(7044): p. 983–987.

41. Lim, K.J.H., et al., Structure-Guided Engineering of Prenyltransferase NphB for High-Yield and Regioselective Cannabinoid Production. ACS Catalysis, 2022. 12(8): p. 4628–4639.

42. Qian, S., J.M. Clomburg, and R. Gonzalez, Engineering Escherichia coli as a platform for the in vivo synthesis of prenylated aromatics. Biotechnology and bioengineering, 2019. 116(5): p. 1116–1127.

43. Valliere, M.A., et al., A cell-free platform for the prenylation of natural products and application to cannabinoid production. Nature communications, 2019. 10(1): p. 1–9.

44. Merchant, S.S., et al., The Chlamydomonas genome reveals the evolution of key animal and plant functions. Science, 2007. 318(5848): p. 245–250.

45. Einhaus, A., et al., Engineering a powerful green cell factory for robust photoautotrophic diterpenoid production. Metabolic Engineering, 2022.

46. Kong, F., et al., Molecular genetic tools and emerging synthetic biology strategies to increase cellular oil content in Chlamydomonas reinhardtii. Plant and Cell Physiology, 2019. 60(6): p. 1184–1196.

47. Schroda, M. and C. Remacle, Molecular advancements establishing Chlamydomonas as a host for biotechnological exploitation. Frontiers in Plant Science, 2022. 13: p. 911483.

48. Schroda, M., Good news for nuclear transgene expression in Chlamydomonas. Cells, 2019. 8(12): p. 1534.

49. Gallaher, S.D., et al., Chlamydomonas genome resource for laboratory strains reveals a mosaic of sequence variation, identifies true strain histories, and enables strain-specific studies. The Plant Cell, 2015. 27(9): p. 2335–2352.

50. Beauchemin, R., et al., Successful reversal of transgene silencing in Chlamydomonas reinhardtii. Biotechnology Journal, 2024. 19(1): p. 2300232.

51. Baier, T., et al., Introns mediate post-transcriptional enhancement of nuclear gene expression in the green microalga Chlamydomonas reinhardtii. PLoS genetics, 2020. 16(7): p. e1008944.

52. Baier, T., et al., Intron-containing algal transgenes mediate efficient recombinant gene expression in the green microalga Chlamydomonas reinhardtii. Nucleic acids research, 2018. 46(13): p. 6909–6919.

53. Barolo, L., et al., Proteomic analysis reveals molecular changes following genetic engineering in Chlamydomonas reinhardtii. BMC Microbiology, 2024. 24(1): p. 392.

54. Zhao, M.-L., et al., Metabolic engineering of the isopentenol utilization pathway enhanced the production of terpenoids in Chlamydomonas reinhardtii. Marine Drugs, 2022. 20(9): p. 577.

55. Lauersen, K.J., et al., Phototrophic production of heterologous diterpenoids and a hydroxy-functionalized derivative from Chlamydomonas reinhardtii. Metabolic engineering, 2018. 49: p. 116–127.

56. Lauersen, K.J., et al., Efficient phototrophic production of a high-value sesquiterpenoid from the eukaryotic microalga Chlamydomonas reinhardtii. Metabolic engineering, 2016. 38: p. 331–343.

57. Gorman, D.S. and R. Levine, Cytochrome f and plastocyanin: their sequence in the photosynthetic electron transport chain of Chlamydomonas reinhardi. Proceedings of the National Academy of Sciences, 1965. 54(6): p. 1665–1669.

58. Kropat, J., et al., A revised mineral nutrient supplement increases biomass and growth rate in Chlamydomonas reinhardtii. The Plant Journal, 2011. 66(5): p. 770–780.

59. Harris, E.H. and E.H. Harrisduke, The Chlamydomonas Sourcebook: Introduction to Chlamydomonas and Its Laboratory Use: Volume 1. 2009: Elsevier Science.

60. Lauersen, K.J., O. Kruse, and J.H. Mussgnug, Targeted expression of nuclear transgenes in Chlamydomonas reinhardtii with a versatile, modular vector toolkit. Applied microbiology and biotechnology, 2015. 99(8): p. 3491–3503.

61. Fischer, N. and J.-D. Rochaix, The flanking regions of PsaD drive efficient gene expression in the nucleus of the green alga Chlamydomonas reinhardtii. Molecular Genetics and Genomics, 2001. 265: p. 888–894.

62. Plucinak, T.M., et al., Improved and versatile viral 2 A platforms for dependable and inducible high level expression of dicistronic nuclear genes in Chlamydomonas reinhardtii. The Plant Journal, 2015. 82(4): p. 717–729.

63. Taura, F., et al., Cannabidiolic-acid synthase, the chemotype-determining enzyme in the fiber-type Cannabis sativa. FEBS letters, 2007. 581(16): p. 2929–2934.

64. Dälken, B., et al., Maltose-Binding Protein Enhances Secretion of Recombinant Human Granzyme B Accompanied by In Vivo Processing of a Precursor MBP Fusion Protein. PLOS ONE, 2010. 5(12): p. e14404.

65. Duong-Ly, K.C. and S.B. Gabelli, Affinity Purification of a Recombinant Protein Expressed as a Fusion with the Maltose-Binding Protein (MBP) Tag. Methods Enzymol, 2015. 559: p. 17–26.

66. Azevedo, F., H. Pereira, and B. Johansson, Colony PCR, in PCR. 2017, Springer. p. 129–139.

67. Nouemssi, S.B., et al., Rapid and efficient colony-PCR for high throughput screening of genetically transformed Chlamydomonas reinhardtii. Life, 2020. 10(9): p. 186.

68. Kindle, K.L., High-frequency nuclear transformation of Chlamydomonas reinhardtii. Proceedings of the National Academy of Sciences, 1990. 87(3): p. 1228–1232.

69. Geissmann, Q., OpenCFU, a new free and open-source software to count cell colonies and other circular objects. PloS one, 2013. 8(2).

70. Neupert, J., et al., An epigenetic gene silencing pathway selectively acting on transgenic DNA in the green alga Chlamydomonas. Nature communications, 2020. 11(1): p. 1–17.

71. Livak, K.J. and T.D. Schmittgen, Analysis of relative gene expression data using real-time quantitative PCR and the 2− ΔΔCT method. methods, 2001. 25(4): p. 402–408.

72. Schroda, M., C.F. Beck, and O. Vallon, Sequence elements within an HSP70 promoter counteract transcriptional transgene silencing in Chlamydomonas. The Plant Journal, 2002. 31(4): p. 445–455.

73. Jinkerson, R.E. and M.C. Jonikas, Molecular techniques to interrogate and edit the Chlamydomonas nuclear genome. The Plant Journal, 2015. 82(3): p. 393–412.

74. Zhang, R., et al., High-throughput genotyping of green algal mutants reveals random distribution of mutagenic insertion sites and endonucleolytic cleavage of transforming DNA. The Plant Cell, 2014. 26(4): p. 1398–1409.

75. Geisler, K., et al., Exploring the impact of terminators on transgene expression in Chlamydomonas reinhardtii with a synthetic biology approach. Life, 2021. 11(9): p. 964.

76. Donnelly, M.L., et al., The cleavage activities of aphthovirus and cardiovirus 2A proteins. Journal of General Virology, 1997. 78(1): p. 13–21.

77. Luke, G.A. and M.D. Ryan, The 2A Story: The End of the Beginning. Beyond the Blueprint-Decoding the Elegance of Gene Expression: Decoding the Elegance of Gene Expression, 2024: p. 115.

78. Zirpel, B., et al., *Optimization of* Δ*9-tetrahydrocannabinolic acid synthase production in Komagataella phaffii via post-translational bottleneck identification*. Journal of biotechnology, 2018. 272: p. 40–47.

79. Geissler, M., et al., *Subcellular localization defines modification and production of* Δ *9-tetrahydrocannabinolic acid synthase in transiently transformed Nicotiana benthamiana*. Biotechnology letters, 2018. 40: p. 981–987.

80. Zirpel, B., F. Stehle, and O. Kayser, *Production of* Δ*9-tetrahydrocannabinolic acid from cannabigerolic acid by whole cells of Pichia (Komagataella) pastoris expressing* Δ*9-tetrahydrocannabinolic acid synthase from Cannabis sativa L*. Biotechnology letters, 2015. 37: p. 1869–1875.

81. Tran, N.T. and R. Kaldenhoff, Achievements and challenges of genetic engineering of the model green alga Chlamydomonas reinhardtii. Algal Research, 2020. 50: p. 101986.

82. Kumar, A., V.R. Falcao, and R.T. Sayre, Evaluating nuclear transgene expression systems in Chlamydomonas reinhardtii. Algal research, 2013. 2(4): p. 321–332.

83. Goold, H.D., J.L. Moseley, and K.J. Lauersen, The synthetic future of algal genomes. Cell Genomics, 2024.

84. Wichmann, J., et al., *Tailored carbon partitioning for phototrophic production of (E)-*α*-bisabolene from the green microalga Chlamydomonas reinhardtii*. Metabolic engineering, 2018. 45: p. 211–222.

85. Mayfield, S.P., et al., Chlamydomonas reinhardtii chloroplasts as protein factories. Current opinion in biotechnology, 2007. 18(2): p. 126–133.

86. Zhang, Z., et al., Efficient heterotrophic cultivation of Chlamydomonas reinhardtii. Journal of Applied Phycology, 2019. 31: p. 1545–1554.

87. Bolaños-Martínez, O.C., et al., Engineering Nicotiana benthamiana for production of active cannabinoid synthases via secretory pathway optimization. Biotechnology Reports, 2025. 45: p. e00865.

88. Barolo, L., et al., Unassembled cell wall proteins form aggregates in the extracellular space of Chlamydomonas reinhardtii strain UVM4. Applied Microbiology and Biotechnology, 2022. 106(11): p. 4145–4156.

89. Ramos Martinez, E.M.L. Fimognari, and Y. Sakuragi, High yield secretion of recombinant proteins from the microalga Chlamydomonas reinhardtii. Plant biotechnology journal, 2017. 15(9): p. 1214–1224.

90. Chung, K.P., et al., Identification and characterization of the COPII vesicle forming GTPase Sar1 in Chlamydomonas. Plant Direct, 2024. 8(6): p. e614.

91. Perozeni, F., et al., Towards microalga-based superfoods: heterologous expression of zeolin in Chlamydomonas reinhardtii. Frontiers in Plant Science, 2023. 14: p. 1184064.

92. Rasala, B.A., et al., Enhanced genetic tools for engineering multigene traits into green algae. PloS one, 2014. 9(4): p. e94028.

93. Dementyeva, P., et al., A novel, robust and mating-competent Chlamydomonas reinhardtii strain with an enhanced transgene expression capacity for algal biotechnology. Biotechnol Rep (Amst), 2021. 31: p. e00644.

94. Shoyama, Y., et al., *Structure and function of*Δ *1-tetrahydrocannabinolic acid (THCA) synthase, the enzyme controlling the psychoactivity of Cannabis sativa*. Journal of molecular biology, 2012. 423(1): p. 96–105.

95. Aschern, M., et al., A novel MoClo-mediated intron insertion system facilitates enhanced transgene expression in Chlamydomonas reinhardtii. Frontiers in Plant Science, 2025. 16: p. 1544873.

96. Jaeger, D., T. Baier, and K.J. Lauersen, Intronserter, an advanced online tool for design of intron containing transgenes. Algal research, 2019. 42: p. 101588.

97. Lumbreras, V., D.R. Stevens, and S. Purton, Efficient foreign gene expression in Chlamydomonas reinhardtii mediated by an endogenous intron. The Plant Journal, 1998. 14(4): p. 441–447.

98. Rose, A.B., Introns as Gene Regulators: A Brick on the Accelerator. Frontiers in Genetics, 2019. 9(672).

99. Einhaus, A., et al., Rational promoter engineering enables robust terpene production in microalgae. ACS Synthetic Biology, 2021. 10(4): p. 847–856.

100. Milito, A., et al., Rational Design and Screening of Synthetic Promoters in Chlamydomonas reinhardtii, in Synthetic Promoters: Methods and Protocols. 2024, Springer. p. 69–83.

101. Milito, A., et al., Challenges and advances towards the rational design of microalgal synthetic promoters in Chlamydomonas reinhardtii. Journal of Experimental Botany, 2023. 74(13): p. 3833–3850.

102. Crozet, P., et al., Birth of a photosynthetic chassis: a MoClo toolkit enabling synthetic biology in the microalga Chlamydomonas reinhardtii. ACS synthetic biology, 2018. 7(9): p. 2074–2086.

103. Ghribi, M., et al., Genome editing by CRISPR-Cas: a game change in the genetic manipulation of Chlamydomonas. Life, 2020. 10(11): p. 295.

104. López Paz, C., et al., Identification of Chlamydomonas reinhardtii endogenous genic flanking sequences for improved transgene expression. The Plant Journal, 2017. 92(6): p. 1232–1244.

105. Mohamadnia, S., et al., Progress and prospects in metabolic engineering approaches for isoprenoid biosynthesis in microalgae. Biotechnology for Biofuels and Bioproducts, 2025. 18(1): p. 64.

106. Einhaus, A., et al., Genome editing of epigenetic transgene silencing in Chlamydomonas reinhardtii. Trends in Biotechnology, 2025.

