## Supporting Information for "Insights into cannabinoid biosynthesis in *Chlamydomonas reinhardtii* : successes with *NphB* and limitations of *CBDAS* expression"

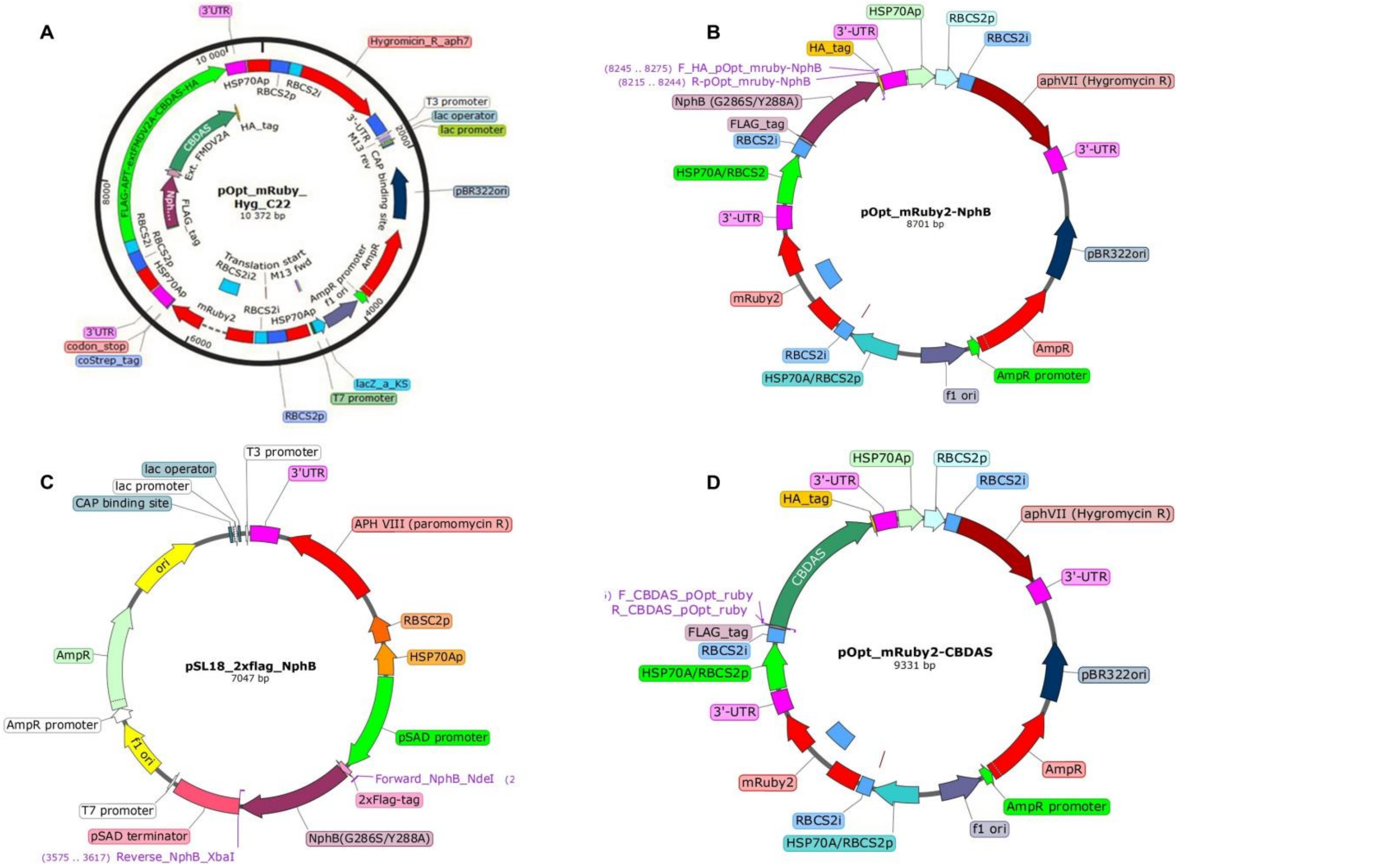

**Figure S1**. Recombinant vectors pOpt_mRuby2-*aph*VII and pSL18-*aph*VIII harboring A) the bicistronic constructs C1, of *NphB*-2A-*CBDAS* under ARp regulation; B) the monocistronic construct C2, of *NphB* under ARp regulation*;* C) the monocistronic construct C3, of *NphB* under pSAD regulation; D) the monocistronic construct C4, of *CBDAS* under ARp regulation. The vector design was created using SnapGene.

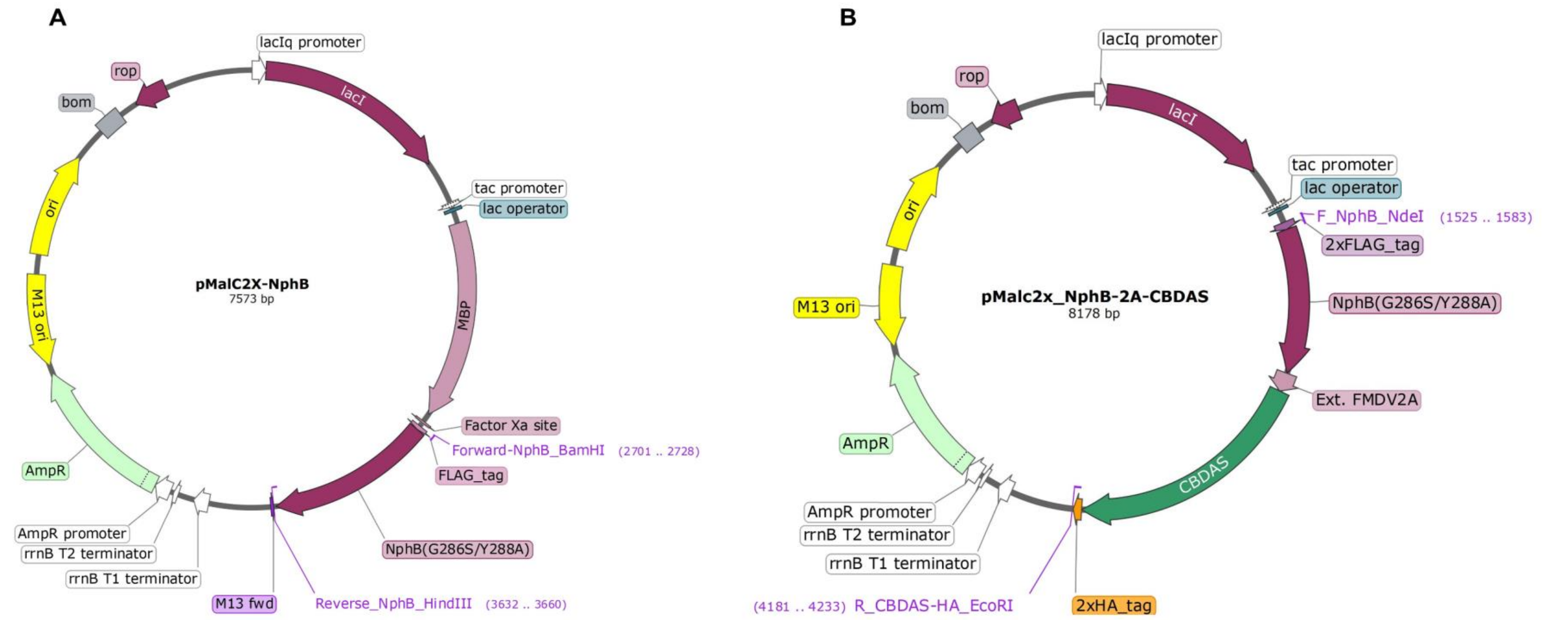

**Figure S2**. Recombinant vectors pMal-c2x harboring NphB and CBDAS genes for expression in *E. coli.* A) pMalc2x holds the monocistronic genetic construct (NphB) fused to the C-terminal region of maltose binding protein (MBP), B) pMalc2x holds the bicistronic genetic construct (NphB-2A-CBDAS) without MBP

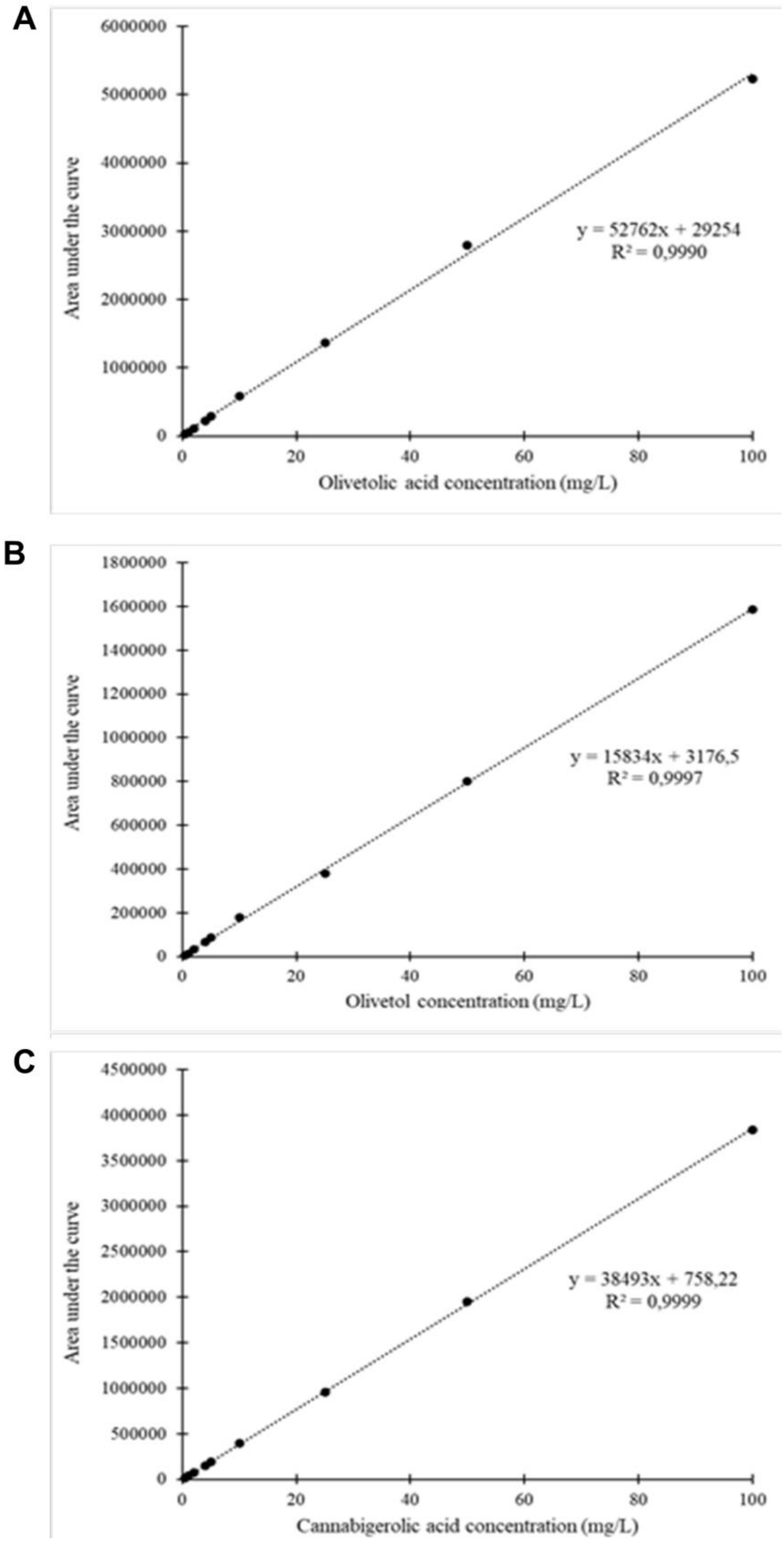

**Figure S3**. Calibration curves. A) Olivetolic acid, B) Olivetol, and C) Cannabigerolic acid.

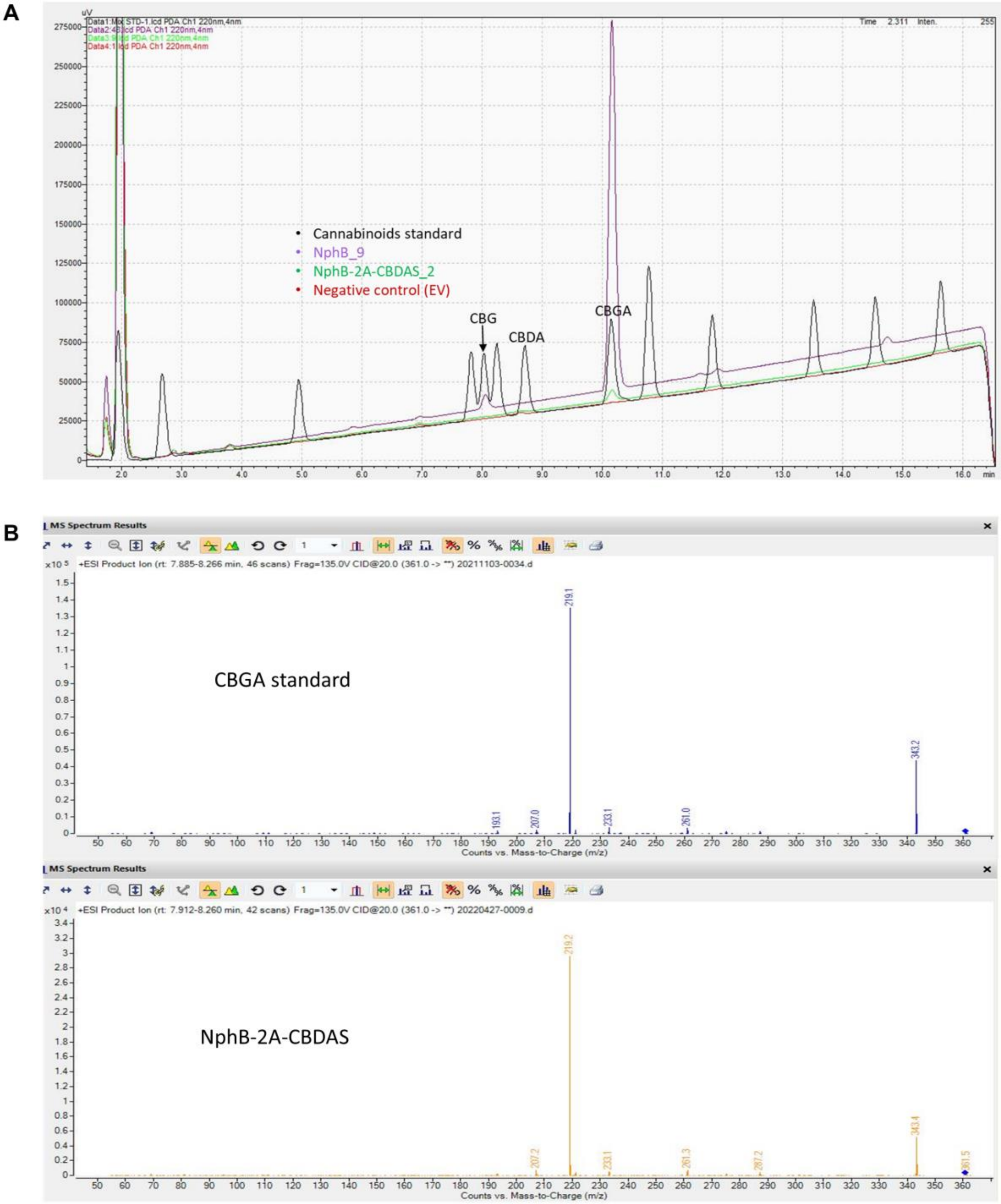

**Figure S4**. Chromatograms showing after *in vitro* enzymatic assay analysis: A) the detection of CBGA at the retention time of 10.146 min when the single protein NphB (peak in purple) or the fused protein NphB-2A-CBDAS (peak in green) was used, with no peak matching the CBDA standard (in black), compared to the empty vector (EV) used (in red); B) Confirmation of CBGA mass-to-charge (m/z) produced *in vitro* using NphB-2A-CBDAS protein compared to the CBGA standard.

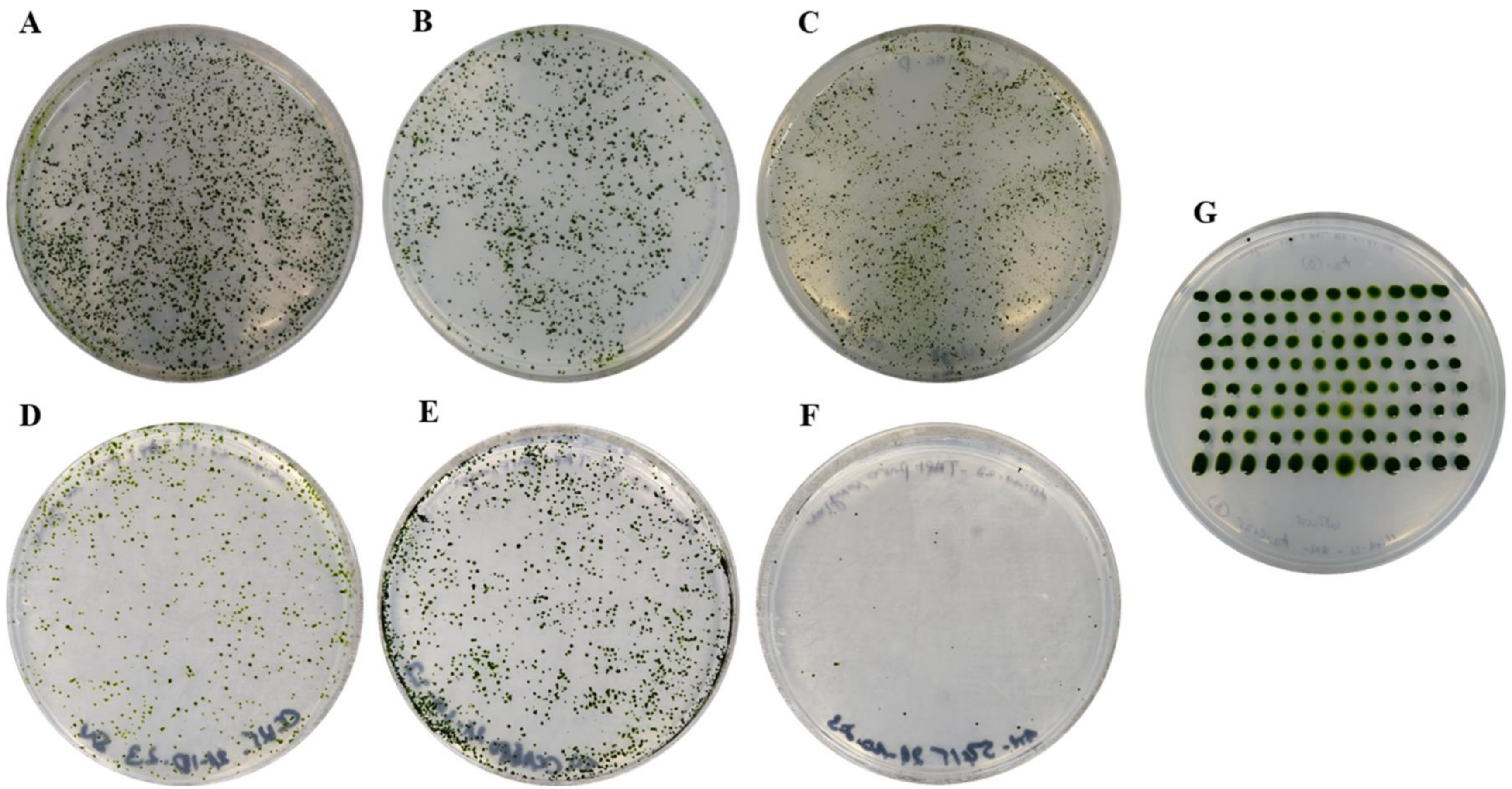

**Figure S5**. Colonies obtained after the nuclear transformation of different *C. reinhardtii* wild-type strains by electroporation with different genetic constructs and a layout of transformants randomly selected on a TAP-hygromycin or paromomycin selection plate. **(A)** Strain CC-125 transformed with C1, **(B)** Strain CC-125 transformed with C2, **(C)** Strain CC-125 co-transformed with C2+C4, **(D)** Strain CC-125 transformed with C3, **(E)** Strain CC-1690 transformed with C3, **(F)** Strain CC-5415 transformed with C3, **(G)** an example of transformants selected growing on TAP media selection plate in a layout of 96 wells plate for high throughput sub-culturing and colony PCR screening. All the strains were transformed with an equal amount of the digested linearized and purified recombinant DNA (2 µg).

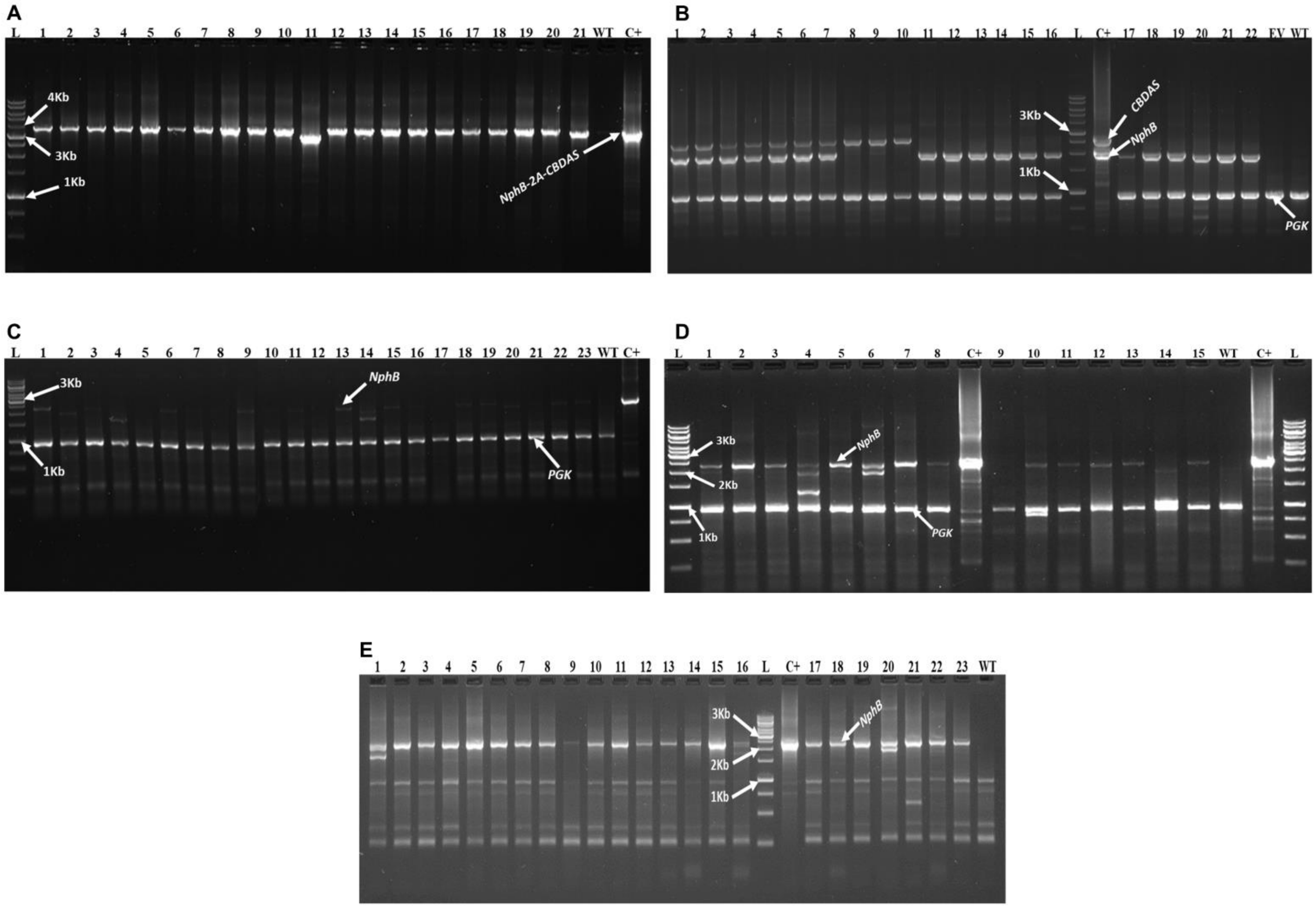

**Figure S6**. Illustration of 1% agarose gels showing transgenes amplicons after colony PCR screening of transformants selected. (A) C1 amplicons of 3.5 kb fromCC-125 transformants, (B) An example of C2 and C4 amplification of 2.577 kb and 1.950 kb amplicons, respectively, from CC-125 transformants, (C) C3 amplicons of 2.363 kb from CC-125 transformants, (D) C3 amplicons of 2.363 kb from CC-5415 transformants, and (E) C3 amplicons of 2.363 kb from CC-1690 transformants. PCR products were run for 45 to 60 min at 100 V constant, and positive control (C+) used was the purified recombinant plasmid DNA of the corresponding construct in each case. The PGK (Phosphoglycerate Kinase: amplicon size 0.944 kb) was used as an internal control for gDNA extraction in the high throughput colony PCR experiments. L: Frogga Bio 1 Kb DNA ladder; WT: wild-type strains.

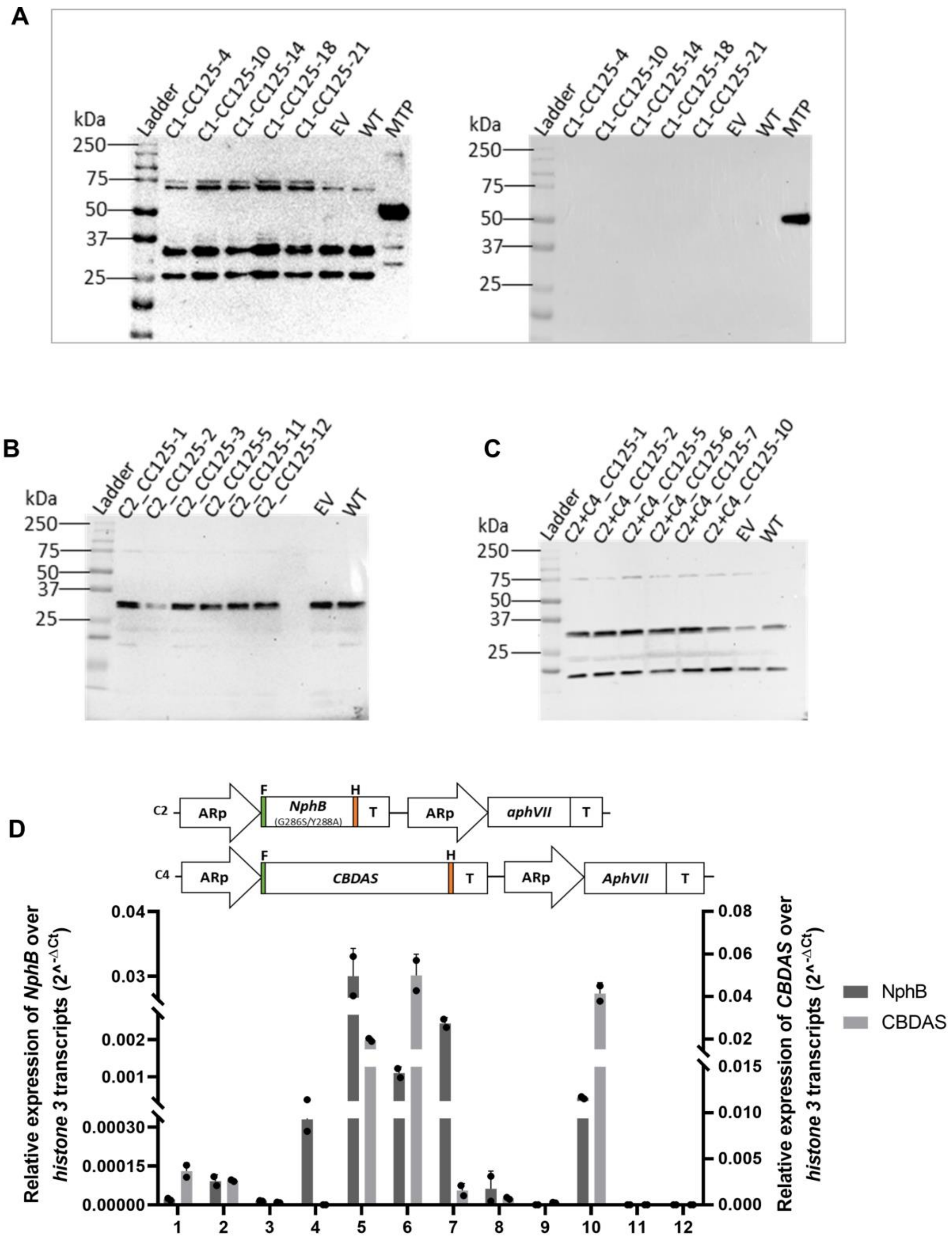

**Figure S7**. *C. reinhardtii* strain CC-125 transformed or co-transformed with the genetic constructs C1, C2, and C4 successfully expressed or co-expressed the transgenes NphB and CBDAS but did not yield the expected proteins. A) Western blot showing unspecific bands detection (left panel) onto the membrane containing separated protein from transformants expressing C1 when treated with Flag antibody; on the right panel is the membrane treated with HA antibody, which shows no band excepted the multi-tag purified protein(MTP) used as a positive control for the experiment; B) Expressed NphB protein extracts after Western blot analysis showing unspecific bands after treatment with flag antibody; C) Co-expressed NphB and CBDAS protein extracts after Western blot analysis showing unspecific bands after treatment with flag antibody; D) NphB and CBDAS expression levels in transformants obtained from the co-transformation of genetic constructs C2 and C4The expected sizes of proteins of interest are C1≈100 kDa or ≈37 kDa and ≈ 60 kDa(if the ribosome-skip mediated by 2A occurs), C2≈ 37 kDa, and C4≈ 60 kDa. EV: empty vector, WT: wild-type. n=2, results are shown as mean ± standard deviation.

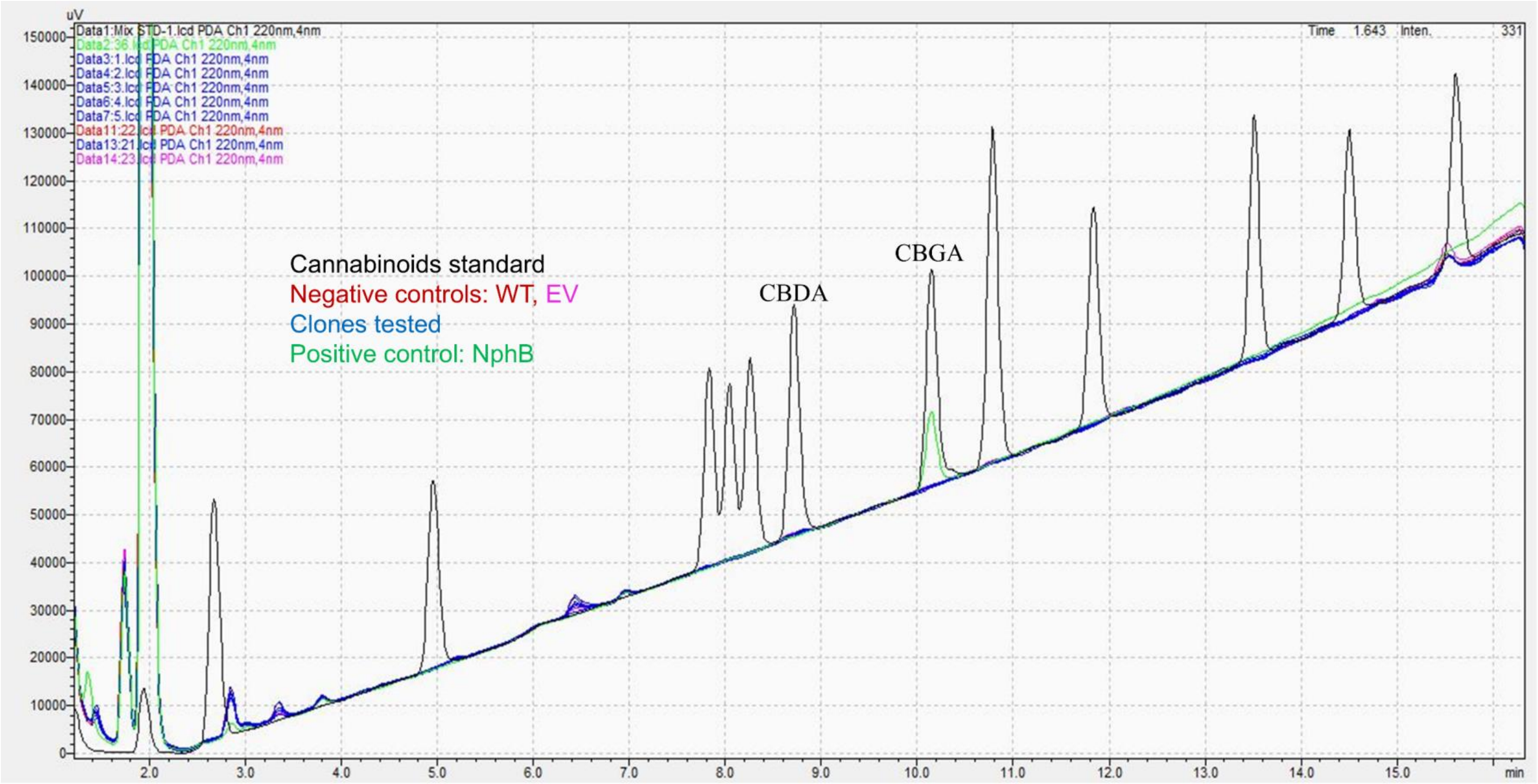

**Figure S8**. HPLC-DAD chromatograms of *in vitro* enzymatic assays with *C. reinhardtii* transformants expressing *NphB-2A-CBDAS*. Chromatograms from tested clones (blue) showed no CBGA or CBDA peaks, similar to negative controls (WT, red; EV, pink). In contrast, the positive control (NphB expressed in *E. coli*, green) produced a CBGA peak at a retention time of 10.146 min, which matched the commercial CBGA standard (black).

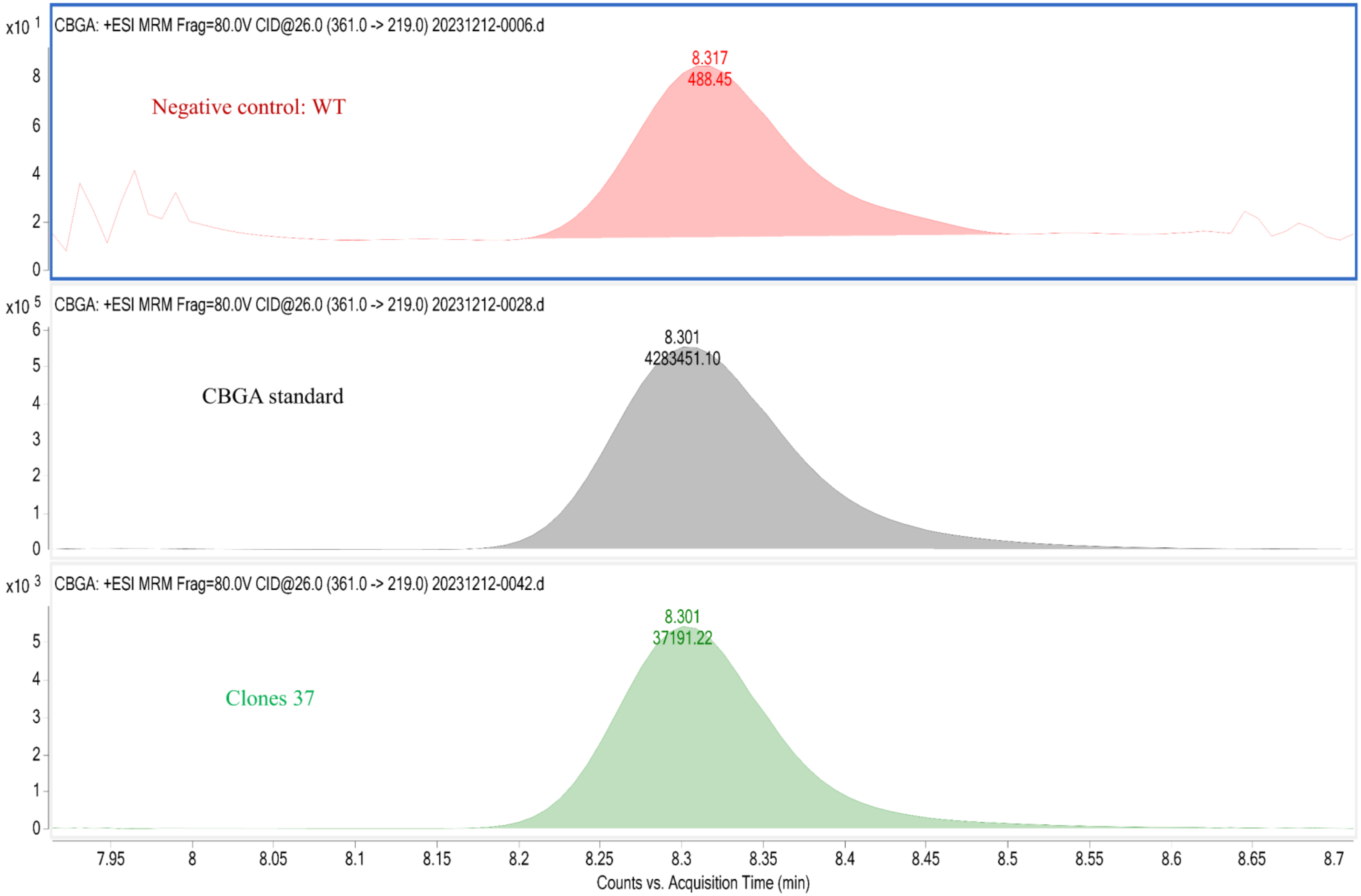

**Figure S9**. CBGA signal detection by HPLC-MS/MS at the retention time of 8.301 min in the positive clone (green) compared to the CBGA standard (black) and the negative control WT (red).

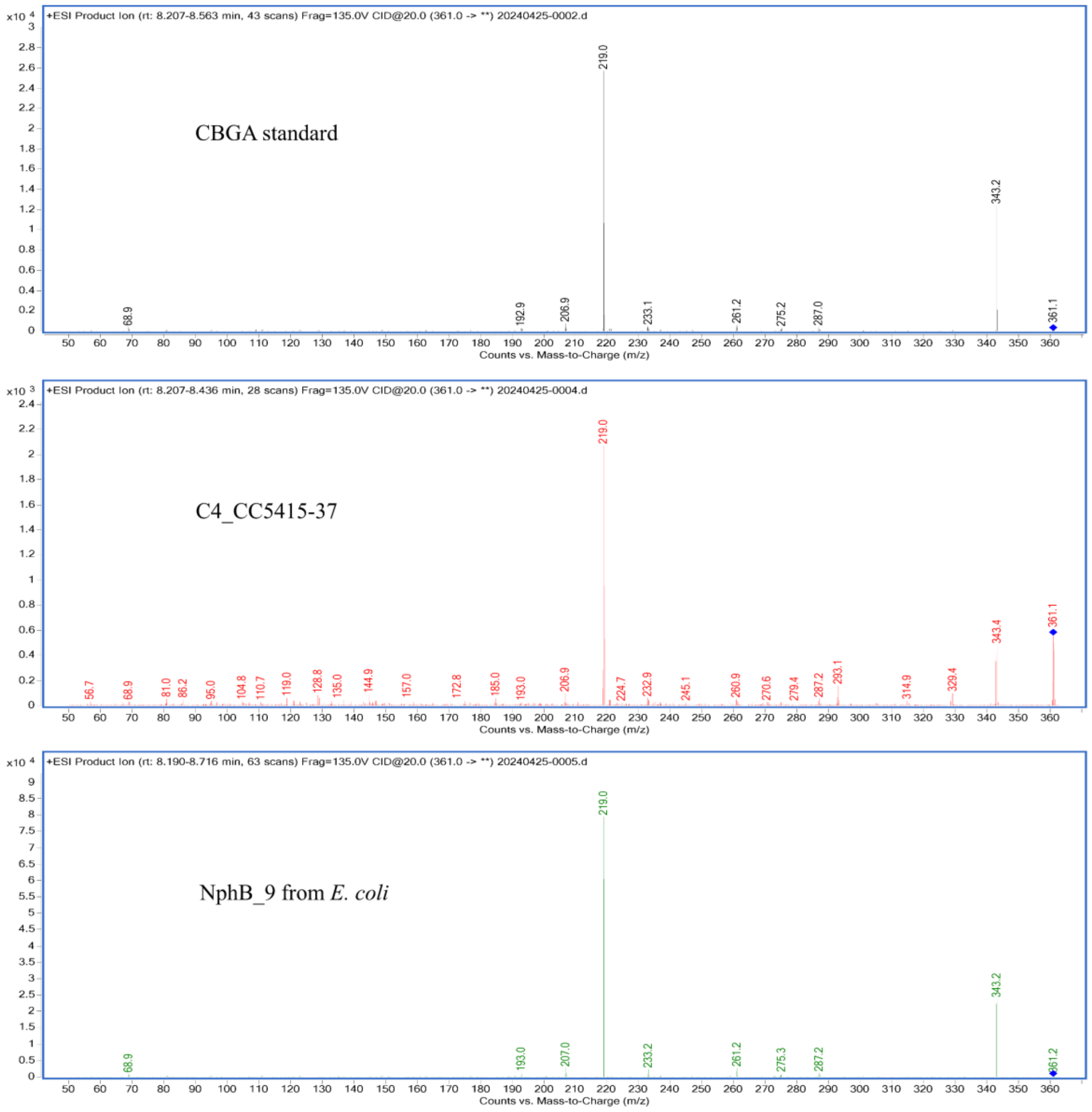

**Figure S10**. Confirmation of CBGA mass-to-charge (m/z) produced *in* vitro by the WB+ transformants of *C. reinhardtii* and *E.coli* expressing a functionally active NphB.

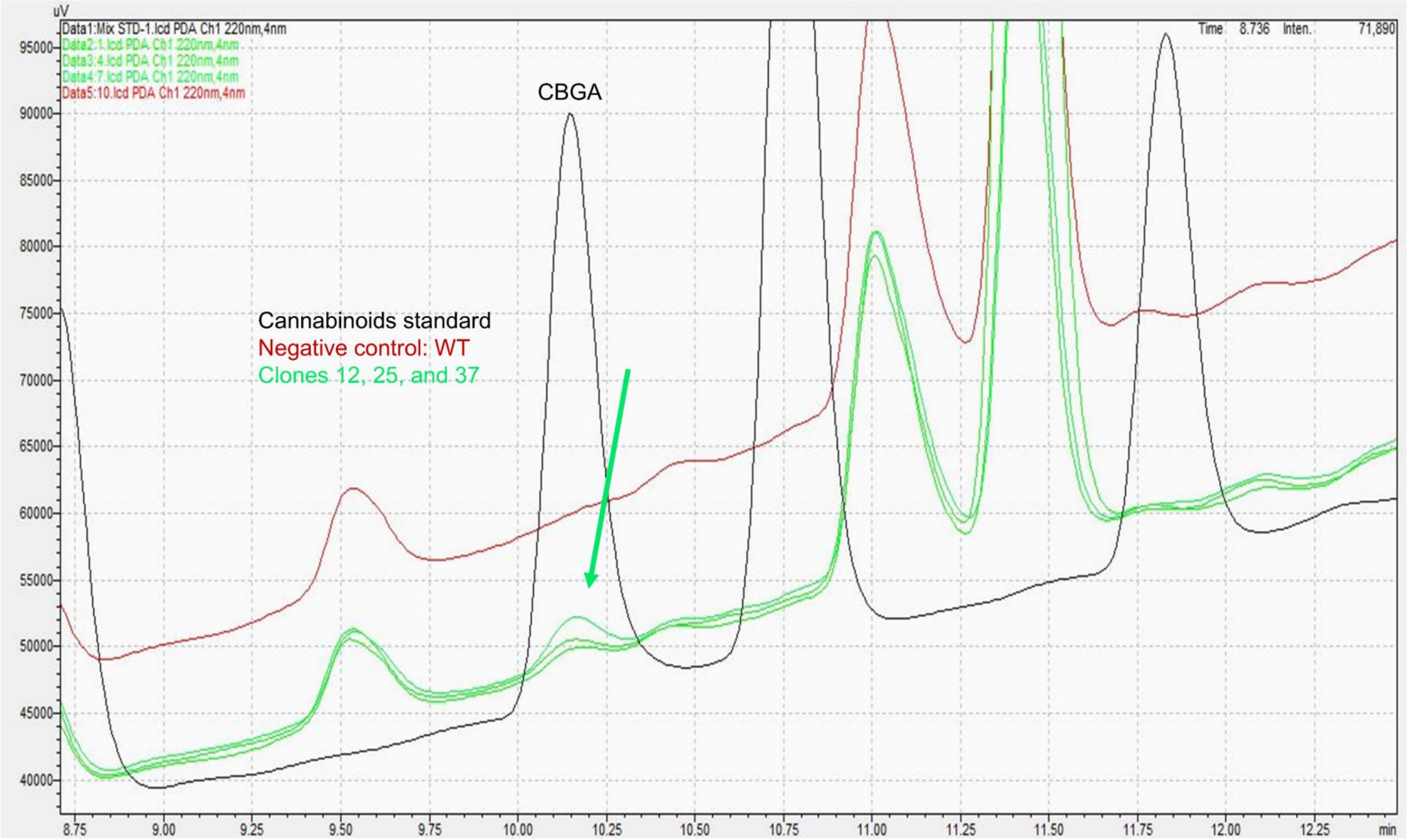

**Figure S11**. Chromatogram showing the CBGA peak at 10.146min (in green highlighted with the arrow) matching the commercial standard CBGA peak (in black) after analysis by HPLC-DAD of *in* vitro enzymatic products obtained from the reaction performed with the *C.reinhardtii* WB+ transformants, compared to the wild-type (WT) in red.

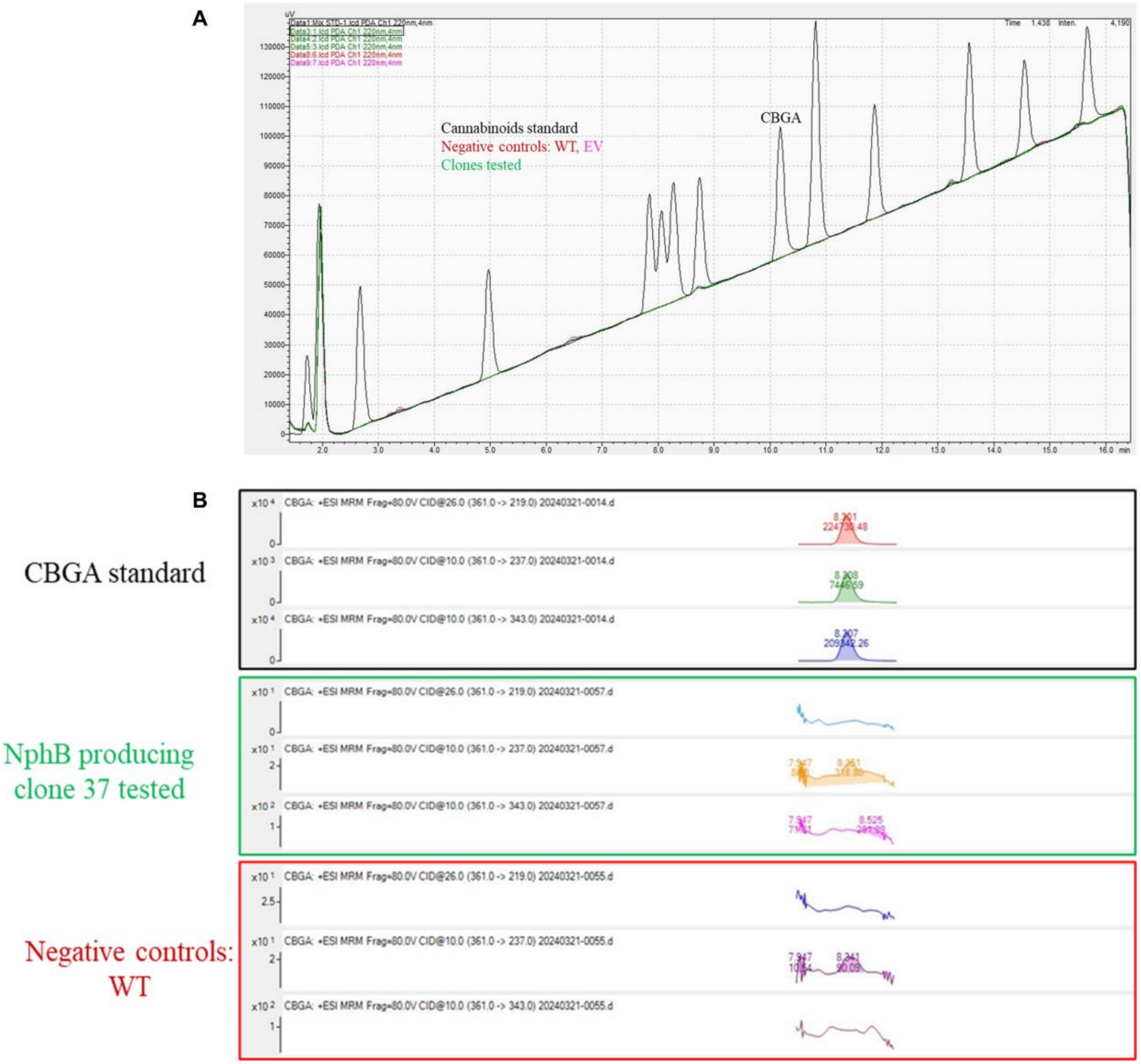

**Figure S12**. HPLC-DAD and HPLC-MS/MS chromatograms, A) showing no detection of CBGA in clones tested (in green), compared to the commercial standard CBGA peak (in black) at the retention time of 10.146 min after analysis by HPLC-DAD of metabolites extracted from clones 12, 25, and 37, respectively, selected from the strains CC-1690, CC-125, and CC-5415; B) the analysis by HPLC-MS/MS further confirmed no detection of CBGA in the clones tested (e.g., clone 37). Chromatograms of clones tested showed a similar profile as the negative controls, wild-type (WT) in red and empty vector (EV) in pink.

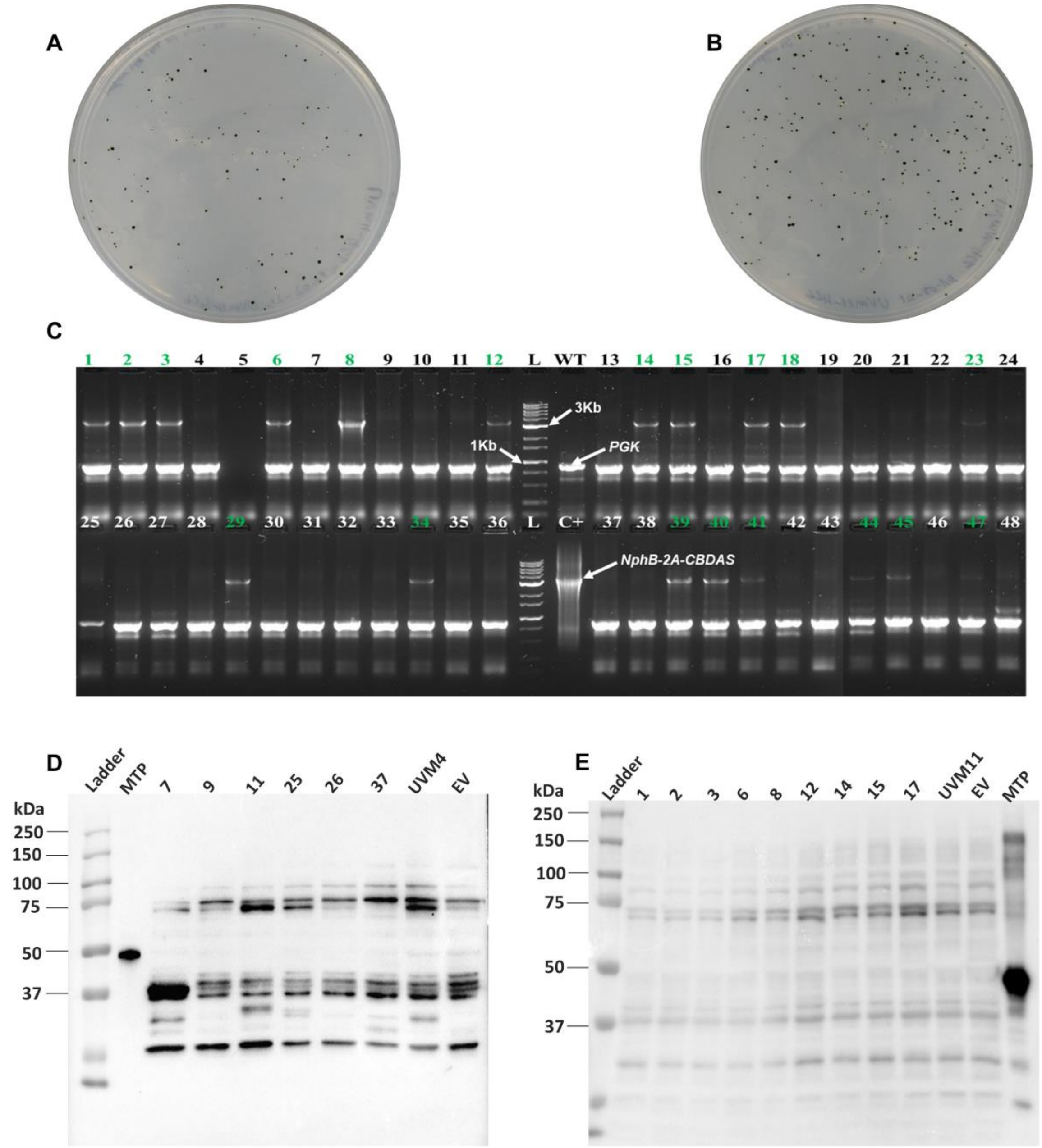

**Figure S13**. Transformation of *C. reinhardtii* strains UVM4 and UVM11 with construct C1. A) Colonies obtained after nuclear transformation of UVM4; B) Colonies obtained after nuclear transformation of UVM11; C) Agarose gel (1%) showing the 3.5 kb transgene amplicon from colony PCR screening of randomly selected UVM11 transformants; D) Western blot of UVM4 transformants expressing C1 probed with anti-FLAG antibody, showing only nonspecific bands; E) Western blot of UVM11 transformants expressing C1 probed with anti-FLAG antibody, also showing nonspecific bands. The expected protein sizes are ~100 kDa for the complete C1 fusion, and ~37 kDa (NphB) and ~60 kDa (CBDAS) if ribosome skipping at the 2A site occurs.

**Supplementary Tables.**

**Table S1.** List of primers used in this study for cloning, colony-PCR, and RT-qPCR

| Genes / constructs | Primer name | Sequence 5'→ 3' |
|---|---|---|
| Cloning into pOPt_mRuby2 (C1, C2, and C4) | | |
| NphB-2A-CBDAS | Ruby2_Hyg_HpaI_C5p1_F | agcgggatccttcgaacgtacggttGCTGAGGCTTGACATGATTGGTG |
| | C22_ScaI_C5p1_R | tgtcgtcgtcgtccttgtagtccatcACTCTGCAAATGGAAACGGCGAC |
| | C5p1_ScaI_C22_F | c gcgtcgccgtttccatttgcagagtgATGGACTACAAGGACGACGACG |
| | C5p2_ScaI_C22_R | cgcctccatttacacggagcggagtCTAGGCGTAGTCAGGCACGTCA |
| | C22_ScaI_C5p2_F | ctatgacgtgcctgactacgcctagACTCCGCTCCGTGTAAATGGAG |
| | Ruby2_Hyg_SnaBI_C5p2_R | gcctcagcacgcgtatttaaattacCGCTTCAAATACGCCCAGCCC |

| | | |
|---|---|---|
| CBDAS | F-pOpt_mRuby-CBDAS | AACCCACGCGAGAACTTCCTGAAGTGCTTCA |
| | R-pOpt_mRuby-CBDAS | CTTGTCGTCGTCGTCCTTGTAGTCCAT |
| NphB | NphB_pOpt_mRuby-F | TACCCCTATGACGTGCCTGACTACGCCTAGACTCC |
| | NphB-pOpt_mRuby-R | GTCCTCCAGGCTGTCGAACGCCTTG |
| | Cloning into pSL18 (C3 and C3b) | |
| NphB | F_NphB_*Nde*I | CGTA**CATATG**ATGGATTACAAGGATGACGACGATAAGGACTACAAGGA CGACGACGACAAGTC |
| | R_NphB_*Xba*I | GTCA**TCTAGA**CTAGTCCTCCAGGCTGTCGAACGCCTTGAGCAGGC |
| CBDAS | F_CBDAS_*Nde*I | **CATATG**ATGAACCCACGCGAGAACTTC |
| | R_CBDAS-HA_*Eco*RI | GCAT**GAATTC**TCAAGCGTAATCTGGAACATCGTATGGGTAGGCGTAGT CAGGCACGT |

| | | |
|---|---|---|
| Cloning into pMAL-c2x and colony-PCR of *E. coli* clones | | |
| NphB-2A-CBDAS | F_NphB_*Nde*I | CGTA**CATATG**ATGGATTACAAGGATGACGACGATAAGGACTACAAGGACGACGACGACAAGTC |
| | R_CBDAS-HA_*EcoR*I | GCAT**GAATTC**TCAAGCGTAATCTGGAACATCGTATGGGTAGGCGTAGTCAGGCACGT |
| NphB | F-NphB/*BamH*I | **GGATCC**ATGTCCGAGGCCGCGG |
| | R-NphB/*Hind*III | **AAGCTT**TTAGTCCTCCAGGCTGTCGAACG |
| Colony PCR of *C. reinhardtii* transformants | | |
| C1, C2, and C4 | Block2_ F | AAGTTCTAGAGTATTTGAAGCGGGATCCTTCGAACGTAC |
| | Block2_R | TACTAAGCTTTGTCAAGCCTCAGCACGCGTATTTAA |
| C3 | F_pSADp-NphB | GCTAGGATCCcacacacctgcccgtctgcctgac |

| | | |
|---|---|---|
| | R_pSADT-NphB | GTCATACGTAcacagtcacgctgtctccccctg |
| RT-qPCR primers were synthesized and validated. | | |
| *PGK* | Cr*PGK*exon9-qP F1 | ATGGGTGTGTTCGAGTTTCC |
| | Cr*PGK*exon9-qP R1 | GTCACCGCCACCAATGAT |
| *Histone* 3 | Cr_*hist*3qPCR FWD Set 2 | GATTGCCCAGGACTTCAAGA |
| | Cr_*hist*3qPCR REV Set 2 | CAGGTTGGTATCCTCGAACAG |
| NphB | Cr_NphB-RT-pcrFor1 | CTGAGCACCTTCCAGGACAC |
| | Cr_NphB-RT-pcrRev1 | GAACAGGCCCTTCTCCACC |
| CBDAS | Cr_CBDAS_RT-pcrFor | TGACGCCCTACGTGAGCAAG |
| | Cr_CBDAS_RT-pcrRev | CCACCAGAGTCTTCACCTTAAC |

**Table S2** Gene sequences used in this study. CDS = coding sequence, SM = selection marker.

| Name | Type | Gene sequence optimized | References |
| --- | --- | --- | --- |
| $NphB_{G286S/Y288A}$ | CDS | ATGTCCGAGGCCGCGGACGTGGAGCGCGTGTACGCGGCCAT<br>GGAGGAGGCCGCCGGCCTGCTGGGCGTGGCCTGCGCCCGCG<br>ACAAGATCTACCCGCTGCTGAGCACCTTCCAGGACACCCTG<br>GTGGAAGGCGGCAGCGTGGTGGTGTTCAGCATGGCCAGCGG<br>CCGCCACAGCACCGAGCTGGACTTCAGCATCAGCGTGCCCA<br>CCTCCCACGGCGACCCCTACGCCACGGTGGTGGAGAAGGGC<br>CTGTTCCCCGCGACCGGCCACCCCGTGGACGACCTGCTGGC<br>GGACACGCAGAAGCACCTGCCGGTGAGCATGTTCGCCATCG<br>ACGGCGAGGTGACGGGCGGCTTCAAGAAGACCTACGCGTTC<br>TTCCCCACCGACAACATGCCCGGCGTGGCCGAGCTCTCGGC<br>GATCCCCTCGATGCCCCCCGCCGTGGCCGAGAACGCGGAGC<br>TGTTCGCGCGGTACGGCCTGGACAAGGTGCAGATGACGTCC<br>ATGGACTACAAGAAGCGCCAGGTGAACCTGTACTTCTCCGA<br>GCTGTCGGCGCAGACCCTGGAGGCCGAGAGCGTGCTGGCCC<br>TGGTGCGGGAGCTGGGCCTGCACGTGCCCAACGAGCTGGGC<br>CTGAAGTTCTGCAAGCGCTCCTTCTCCGTGTACCCCACCCTG<br>AACTGGGAGACCGGCAAGATTGACCGCCTGTGCTTCGCCGT<br>GATTAGCAACGACCCCACCCTGGTGCCCTCCAGCGACGAGG<br>GCGACATCGAGAAGTTCCACAACTACGCCACCAAGGCGCCG<br>TACGCCTACGTGGGCGAGAAGCGCACCCTGGTGTACGGCCT<br>TACCCTGAGCCCCAAGGAGGAGTATTACAAGCTGTCGGCCG<br>CCTATCACATCACCGACGTCCAGCGCGGCCTGCTCAAGGCG<br>TTCGACAGCCTGGAGGACTAG | [1] |
| ExtFMDV2A | CDS | CTGCTGGCCATCCACCCCACCGAGGCCCGGCACAAGCAGAA<br>GATCGTGGCCCCCGTCAAGCAGACGCTGAACTTCGACCTGC<br>TGAAGCTGGCCGGCGACGTGGAGTCGAACCCCGGCCCC | [2] |

| | | | |
|---|---|---|---|
| CBDAS | CDS | ATGAACCCACGCGAGAACTTCCTGAAGTGCTTCAGCCAGTA<br>CATCCCGAACAACGCCACCAACCTGAAGCTGGTCTATACCC<br>AGAACAACCCCCTGTACATGTCCGTCCTGAACAGCACCATC<br>CACAACCTGCGCTTCACCAGCGACACGACCCCGAAGCCCCT<br>CGTGATCGTGACCCCCAGCCACGTGTCCCACATCCAGGGCA<br>CCATCCTGTGCTCGAAGAAGGTGGGCCTGCAGATCCGCACC<br>CGCAGCGGCGGCCACGACTCTGAGGGCATGTCGTACATCAG<br>CCAGGTGCCGTTCGTGATCGTCGACCTGCGCAACATGCGCTC<br>CATCAAGATCGACGTGCACTCGCAGACCGCCTGGGTGGAGG<br>CGGGGGCCACCCTCGGCGAGGTCTACTACTGGGTGAACGAG<br>AAGAACGAGAACCTGAGCCTGGCCGCCGGCTACTGCCCGAC<br>GGTCTGCGCGGGCGGCCACTTCGGCGGCGGCGGCTACGGCC<br>CCCTGATGCGCAACTACGGCCTGGCGGCCGACAACATCATC<br>GACGCGCACCTCGTGAACGTGCACGGCAAGGTGCTGGACCG<br>CAAGTCGATGGGGGAGGACCTGTTCTGGGCCCTGCGCGGCG<br>GCGGCGCCGAGAGCTTCGGCATCATCGTCGCCTGGAAGATC<br>CGCCTGGTGGCGGTGCCCAAGTCCACCATGTTCAGCGTGAA<br>GAAGATCATGGAGATCCACGAGCTCGTGAAGCTGGTCAACA<br>AGTGGCAGAACATCGCGTACAAGTACGACAAGGACCTGCTG<br>CTGATGACCCACTTCATCACGCGCAACATCACGGACAACCA<br>GGGCAAGAACAAGACCGCCATCCACACCTACTTCAGCAGCG<br>TGTTCCTGGGCGGCGTGGACAGCCTGGTCGACCTGATGAAC<br>AAGAGCTTCCCCGAGCTGGGCATCAAGAAGACCGACTGCCG<br>CCAGCTGTCGTGGATCGACACCATCATCTTCTACTCCGGCGT<br>CGTGAACTACGACACCGACAACTTCAACAAGGAGATCCTGC<br>TGGACCGCTCCGCGGGCCAGAACGGCGCGTTCAAGATCAAG<br>CTGGACTACGTGAAGAAGCCCATCCCGGAGAGCGTGTTCGT<br>GCAGATCCTGGAGAAGCTGTACGAGGAGGACATTGGCGCCG<br>GCATGTACGCGCTGTACCCCTACGGCGGCATCATGGACGAG<br>ATCTCCGAGTCCGCCATCCCGTTCCCCCACCGGGCGGGCATC<br>CTGTACGAGCTGTGGTACATCTGCAGCTGGGAGAAGCAGGA | [3] |

| | | | |
|---|---|---|---|
| | | GGACAACGAGAAGCACCTGAACTGGATCCGCAACATCTACA ACTTCATGACGCCCTACGTGAGCAAGAACCCCCGCCTGGCC TACCTGAACTACCGCGACCTGGACATCGGCATCAACGACCC CAAGAACCCGAACAACTACACGCAGGCCCGCATTTGGGGCG AGAAGTATTTCGGCAAGAACTTCGACCGATTGGTTAAGGTG AAGACTCTGGTGGATCCTAATAACTTCTTCCGTAATGAGCAG TCCATCCCCCCCCTGCCGCGGCACCGCCACTAG | |
| Hsp70A-<br>RBCS2/5'UTR | Promoter | GCTGAGGCTTGACATGATTGGTGCGTATGTTTGTATGAAGCT ACAGGACTGATTTGGCGGGCTATGAGGGCGGGGGAAGCTCT GGAAGGGCCGCGATGGGGCGCGCGGCGTCCAGAAGGCGCC ATACGGCCCGCTGGCGGCACCCATCCGGTATAAAAGCCCGC GACCCCGAACGGTGACCTCCACTTTCAGCGACAAACGAGCA CTTATACATACGCGACTATTCTGCCGCTATACATAACCACTC AGCTAGCTTAAGATCCCATCACCGGTGCATGCCGGGCGCGC CAGAAGGAGCGCAGCCAAACCAGGATGATGTTTGATGGGGT ATTTGAGCACTTGCAACCCTTATCCGGAAGCCCCCTGGCCCA CAAAGGCTAGGCGCCAATGCAAGCAGTTCGCATGCAGCCCC TGGAGCGGTGCCCTCCTGATAAACCGGCCAGGGGGCCTATG TTCTTTACTTTTTTACAAGAGAAGTCACTCAACATCTTAAAA TG | [4, 5] |
| RBCS2/3'UTR | Terminator | ACTCCGCTCCGTGTAAATGGAGGCGCTCGTTGATCTGAGCCT TGCCCCCTGACGAACGGCGGTGGATGGAAGATACTGCTCTC AAGTGCTGAAGCGGTAGCTTAGCTCCCCGTTTCGTGCTGATC AGTCTTTTTCAACACGTAAAAAGCGGAGGAGTTTTGCAATTT TGTTGGTTGTAACGATCCTCCGTTGATTTTGGCCTCTTTCTCC ATGGGCGGGCTGGGCGTATTTGAAGCG | |
| PSADp | Promoter | CACACACCTGCCCGTCTGCCTGACAGGAAGTGAACGCATGT CGAGGGAGGCCTCACCAATCGTCACACGAGCCCTCGTCAGA AACACGTCTCCGCCACGCTCTCCCTCTCACGGCCGACCCCGC AGCCCTTTTGCCCTTTCCTAGGCCACCGACAGGACCCAGGCG CTCTCAGCATGCCTCAACAACCCGTACTCGTGCCAGCGGTGC | [6] |

| | | | |
|---|---|---|---|
| | | CCTTGTGCTGGTGATCGCTTGGAAGCGCATGCGAAGACGAA<br>GGGGCGGAGCAGGCGGCCTGGCTGTTCGAAGGGCTCGCCGC<br>CAGTTCGGGTGCCTTTCTCCACGCGCGCCTCCACACCTACCG<br>ATGCGTGAAGGCAGGCAAATGCTCATGTTTGCCCGAACTCG<br>GAGTCCTTAAAAAGCCGCTTCTTGTCGTCGTTCCGAGACATG<br>TTAGCAGATCGCAGTGCCACCTTTCCTGACGCGCTCGGCCCC<br>ATATTCGGACGCAATTGTCATTTGTAGCACAATTGGAGCAA<br>ATCTGGCGAGGCAGTAGGCTTTTAAGTTGCAAGGCGAGAGA<br>GCAAAGTGGGACGCGGCGTGATTATTGGTATTTACGCGACG<br>GCCCGGCGCGTTAGCGGCCCTTCCCCCAGGCCAGGGACGAT<br>TATGTATCAATATTGTTGCGTTCGGGCACTCGTGCGAGGGCT<br>CCTGCGGGCTGGGGAGGGGGATCTGGGAATTGGAGGTACGA<br>CCGAGATGGCTTGCTCGGGGGGAGGTTTCCTCGCCGAGCAA<br>GCCAGGGTTAGGTGTTGCGCTCTTGACTCGTTGTGCATTCTA<br>GGACCCCACTGCTACTCACAACAAGC | |
| PSADt | Terminator | TGGCAGCAGCTGGACCGCCTGTACCATGGAGAAGAGCTTTA<br>CTTGCCGGGATGGCCGATTTCGCTGATTGATACGGGATCGG<br>AGCTCGGAGGCTTTCGCGCTAGGGGCTAGGCGAAGGGCAGT<br>GGTGACCAGGGTCGGTGTGGGGTCGGCCCACGGTCAATTAG<br>CCACAGGAGGATCAGGGGGAGGTAGGCACGTCGACTTGGTT<br>TGCGACCCCGCAGTTTTGGCGGACGTGCTGTTGTAGATGTTA<br>GCGTGTGCGTGAGCCAGTGGCCAACGTGCCACACCCATTGA<br>GAAGACCAACCAACTTACTGGCAATATCTGCCAATGCCATA<br>CTGCATGTAATGGCCAGGCCATGTGAGAGTTTGCCGTGCCT<br>GCGCGCGCCCCGGGGGCGCAGTTTAGCTGACCAGCCGTGGG<br>ATGATGCACGCATTTGCAAGGACAGGGTAATCACAGCAGCA<br>ACATGGTGGGCTTAGGACAGCTGTGGGTCAGTGGACGGACG<br>GCAGGGGAGGGACGGCGCAGCTCGGGAGACAGGGGGAGAC<br>AGCGTGACTGTG | [6] |
| *aphVII* | Hygromycin SM | ATGACACAAGAATCCCTGTTACTTCTCGACCGTATTGATTCG<br>GATGATTCCTACGCGAGCCTGCGGAACGACCAGGAGTTCTG | [4] |

| | | | |
|---|---|---|---|
| | | GGAGCCGCTGGCCCGCCGAGCCCTGGAGGAGCTCGGGCTGC<br>CGGTGCCGCCGGTGCTGCGGGTGCCCGGCGAGAGCACCAAC<br>CCCGTACTGGTCGGCGAGCCCGGCCCGGTGATCAAGCTGTT<br>CGGCGAGCACTGGTGCGGTCCGGAGAGCCTCGCGTCGGAGT<br>CGGAGGCCTACGCGGTCCTGGCGGACGCCCCGGTGCCGGTG<br>CCCCGCCTCCTCGGCCGCGGCGAGCTGCGGCCCGGCACCGG<br>AGCCTGGCCGTGGCCCTACCTGGTGATGAGCCGGATGACCG<br>GCACCACCTGGCGGTCCGCGATGGACGGCACGACCGACCGG<br>AACGCGCTGCTCGCCCTGGCCCGCGAACTCGGCCGGGTGCT<br>CGGCCGGCTGCACAGGGTGCCGCTGACCGGGAACACCGTGC<br>TCACCCCCCATTCCGAGGTCTTCCCGGAACTGCTGCGGGAAC<br>GCCGCGCGGCGACCGTCGAGGACCACCGCGGGTGGGGCTAC<br>CTCTCGCCCCGGCTGCTGGACCGCCTGGAGGACTGGCTGCC<br>GGACGTGGACACGCTGCTGGCCGGCCGCGAACCCCGGTTCG<br>TCCACGGCGACCTGCACGGGACCAACATCTTCGTGGACCTG<br>GCCGCGACCGAGGTCACCGGGATCGTCGACTTCACCGACGT<br>GTATGCGGGAGACTCCCGCTACAGCCTGGTGCAACTGCATC<br>TCAACGCCTTCCGGGGCGACCGCGAGATCCTGGCCGCGCTG<br>CTCGACGGGGCGCAGTGGAAGCGGACCGAGGACTTCGCCCG<br>CGAACTGCTCGCCTTCACCTTCCTGCACGACTTCGAGGTGTT<br>CGAGGAGACCCCGCTGGATCTCTCCGGCTTCACCGATCCGG<br>AGGAACTGGCGCAGTTCCTCTGGGGGCCGCCGGACACCGCC<br>CCCGGCGCCTGA | |
| *aphVIII* | Paromomycin<br><br>SM | TCAGAAGAACTCGTCCAACAGCCGGTAAAACGCCAGCTTTT<br>CCTCCGATACCGCCCCATCCCACCCGCGCCCGTACTCCCGCA<br>GGAACGCCGCGGAACACTCCGGCCCGAACCACGGGTCCTCC<br>TCGTGGGCCAGCTCGCGCAGCACCAGCGCGAGATCGGAGTG<br>CCGGTCCGCACGGCCGACCCGCCCCACGTCGATCAGCCCGG<br>TCACCTCGCAGGTACGAGGGTCGAGCAGCACGTTGTCCGGG<br>CACAGGTGACCGTGGCAAACCGCCAGATCCTCGTCCGCAGG<br>CCGAGTCCGCTCCAGCTCGGCGAGAAGCCGCTCCCCCGACC | [6] |

| | | |
|---|---|---|
| | | ACCCCTTCCGCTCCTCGTCCAGATCCTCCAAGTCGACGCTCC<br>CTTCAGCGACAGCACGGGCCGCCTGCGGCACCGTCACCGCG<br>AGACTGCGATCGAACGGACACCGCTCCCAGTCCAGCGCGTG<br>CAGCGAACGAGCGAGCCCCGCGAGCGCCACCGCCACGTCCA<br>GCCGCTGCTCCCGCGGCCACCGCGCACTGGCCGGACGCCCC<br>GGAACCGCTTCGGTGACCAACCAGGCGACCCTCTCGTCCCC<br>ACCACCCTCCACAACACGAGGTACGGGAATCCCCACCTCCG<br>CCAACCACACCAGCCGCTCAGCCTCACCCAACAAGCCCACC<br>CCGGCCCCCAGAGCTGCCACCTTGACAAACAACTCCCGCCC<br>ACCACCCCGAAGCCGATAAACACCAGCCCCCGAGGCCCCAT<br>CCTCCACAACAACCCACTCACAACCGGGATACCGACCCCGC<br>AGTGCACGCAACGCATCGTCCAT |

**Table S3**. Analytical parameters used for compound identification using high-performance liquid chromatography with diode-array detection (HPLC-DAD). Abbreviations: olivetolic acid (OA), olivetol (OL), delta-9-tetrahydrocannabinol (THC), cannabidiol (CBD), cannabinol (CBN), delta-9-tetrahydrocannabinolic acid (THCA), cannabidiolic acid (CBDA), cannabinolic acid (CBNA), cannabigerolic acid (CBGA), cannabichromene (CBC), cannabigerol (CBG), tetrahydrocannabivarin (THCV), cannabidivarin (CBDV), retention time (RT), maximum absorption wavelengths ($\lambda$max).

| **Compound** | **RT (min)** | **$\lambda$max (nm)** |
|---|---|---|
| OA | 1.936 | 214/261/300 |
| OL | 1.716 | 202/276 |
| THC | 11.827 | 199/279/328 |
| CBD | 7.811 | 199/276 |
| CBN | 10.773 | 198/221/283 |
| THCA | 15.636 | 199/223/270/306 |
| CBDA | 8.709 | 198/223/269/307 |
| CBNA | 14.547 | 199/222/261/328 |
| CBGA | 10.146 | 198/222/268/305 |
| CBC | 13.517 | 199/226/280 |
| CBG | 8.025 | 199/274 |
| THCV | 8.239 | 199/278 |
| CBDV | 4.942 | 200/275 |

**Table S4**. Optimized instrumental parameters used for HPLC-MS/MS analyses in ESI+. References: olivetolic acid (OA), olivetol (OL), delta-9-tetrahydrocannabinol (THC), cannabidiol (CBD), cannabinol (CBN), delta-9-tetrahydrocannabinolic acid (THCA), cannabidiolic acid (CBDA), cannabinolic acid (CBNA), cannabigerolic acid (CBGA), cannabichromene (CBC), cannabigerol (CBG), tetrahydrocannabivarin (THCV), cannabidivarin (CBDV), retention time (RT), collision energy (CE). Quantification MRM transitions are bold, while qualifier MRM transitions are not.

| **Compound** | **RT (min)** | **Parent ion (m/z)** | **Product ion (m/z)** | **Fragmentor (V)** | **CE (V)** | **Polarity** |
|---|---|---|---|---|---|---|
| OA | 4.272 | **225** | **207** | **75** | **10** | **+** |
| | | | 123 | 75 | 18 | + |
| | | | 189 | 75 | 18 | + |
| OL | 4.083 | **181** | **111** | **75** | **10** | **+** |
| | | | 71 | 75 | 10 | + |
| | | | 93 | 75 | 26 | + |
| THC | 9.225 | **315** | **193** | **80** | **22** | **+** |
| | | | 123 | 80 | 34 | + |
| | | | 135 | 80 | 18 | + |
| CBD | 7.752 | **315** | **193** | **75** | **22** | **+** |
| | | | 123 | 75 | 34 | + |
| | | | 135 | 75 | 18 | + |
| CBN | 8.825 | **311** | **223** | **75** | **22** | **+** |
| | | | 293 | 75 | 14 | + |
| | | | 241 | 75 | 18 | + |
| THCA | 10.334 | **359** | **341** | **80** | **14** | **+** |
| | | | 219 | 80 | 34 | + |
| | | | 285 | 80 | 26 | + |

| | | | | | | |
|---|---|---|---|---|---|---|
| CBDA | 7.935 | **359** | **341** | **80** | **10** | **+** |
| | | | 219 | 80 | 30 | + |
| | | | 261 | 80 | 26 | + |
| CBNA | 9.933 | **355** | **337** | **85** | **14** | **+** |
| | | | 253 | 85 | 30 | + |
| | | | 235 | 85 | 30 | + |
| CBGA | 8.289 | **361** | **219** | **80** | **26** | **+** |
| | | | 343 | 80 | 10 | + |
| | | | 237 | 80 | 10 | + |
| CBC | 9.556 | **315** | **193** | **75** | **18** | **+** |
| | | | 259 | 75 | 10 | + |
| | | | 81 | 75 | 10 | + |
| CBG | 7.655 | **317** | **193** | **80** | **14** | **+** |
| | | | 123 | 80 | 34 | + |
| | | | 207 | 80 | 10 | + |
| THCV | 8.162 | **287** | **165** | **75** | **22** | **+** |
| | | | 123 | 75 | 34 | + |
| | | | 231 | 75 | 18 | + |
| CBDV | 6.642 | **287** | **165** | **75** | **22** | **+** |
| | | | 123 | 75 | 34 | + |
| | | | 231 | 75 | 14 | + |

**Table S5.** CBGA peak areas were obtained with the $WB^{+}$ clones from the strains CC-5415, CC-125, and CC-1690 after *in* vitro enzymatic assay analysis by HPLC-MS/MS. ND = Not detected

| *C. reinhardtii* strains | WB+ clones tested | CBGA peak area |
|---|---|---|
| CC-5415 | 4 | 9717.74851 |
| | 5 | 32943.8604 |
| | 6 | 30127.02 |
| | 8 | 8368.85007 |
| | 10 | 16762.1016 |
| | 37 | 34711.9617 |
| | WT | ND |
| | C+ (NphB expressed in Bacteria) | 2904845.28 |
| CC-125 | 25 | 36709,52926 |
| | 29 | 19858.80924 |
| | 64 | 13792.38187 |
| | 66 | 11045.42278 |
| | 85 | 18800.53448 |
| | 95 | 12722.9058 |
| | 97 | 10217.03439 |
| | 98 | ND |
| | 101 | 10856.97318 |
| | 117 | 6758.663872 |
| | 142 | 16607.58164 |
| | 148 | 12622.62841 |
| | 181 | ND |
| | 182 | 4904.456476 |
| | 187 | 6132.950473 |
| | WT | ND |
| | C+ (NphB expressed in Bacteria) | 3069991.921 |

| | | |
|---|---|---|
| CC-1690 | 8 | 30168.62603 |
| | 9 | 8615.292705 |
| | 12 | 83915.14025 |
| | 15 | 46165.52638 |
| | 27 | 81865.52452 |
| | 30 | 56723.32368 |
| | 46 | 44445.57859 |
| | 47 | 5589.331025 |
| | 48 | 7271.732942 |
| | 49 | 4206.08167 |
| | 56 | 21056.19999 |
| | 60 | 13204.18278 |
| | 61 | 5732.458053 |
| | 63 | 10129.23354 |
| | 65 | 2786.874686 |
| | 66 | 30170.85247 |
| | 72 | 7182.685974 |
| | 78 | 27178.03189 |
| | 80 | 8872.063057 |
| | 81 | ND |
| | WT | ND |
| | C+ (NphB expressed in Bacteria) | 3069991.921 |

**Table S6.** Summary of selected UVM4 and UVM11 transformants tested for C1 construct integration and protein accumulation.

| **Genetic construct** | ***C. reinhardtii*** **strains** | **Number of colonies after transformation** | **Randomly selected clones** | **Stable clones after five rounds of subculturing** | **PCR+ clones (%)** | **Clones tested for protein detection** | **WB+ clones (%)** |
|---|---|---|---|---|---|---|---|
| **C1** | UVM4 | 100 | 96 | 91 | 18 (19.8%) | 18 | 0 (0%) |
| | UVM11 | 198 | 96 | 93 | 19 (20.4%) | 19 | 0 (0%) |

**REFERENCES**

1. Valliere, M.A., et al., *A cell-free platform for the prenylation of natural products and application to cannabinoid production.* Nature communications, 2019. **10**(1): p. 1-9.
2. Plucinak, T.M., et al., *Improved and versatile viral 2 A platforms for dependable and inducible high-level expression of dicistronic nuclear genes in C hlamydomonas reinhardtii.* The Plant Journal, 2015. **82**(4): p. 717-729.
3. Luo, X., et al., *Complete biosynthesis of cannabinoids and their unnatural analogues in yeast.* Nature, 2019. **567**(7746): p. 123-126.
4. Lauersen, K.J., O. Kruse, and J.H. Mussgnug, *Targeted expression of nuclear transgenes in Chlamydomonas reinhardtii with a versatile, modular vector toolkit.* Applied microbiology and biotechnology, 2015. **99**(8): p. 3491-3503.
5. Schroda, M., D. Blöcker, and C.F. Beck, *The HSP70A promoter as a tool for the improved expression of transgenes in Chlamydomonas.* The plant journal, 2000. **21**(2): p. 121-131.
6. Fischer, N. and J.-D. Rochaix, *The flanking regions of PsaD drive efficient gene expression in the nucleus of the green alga Chlamydomonas reinhardtii.* Molecular Genetics and Genomics, 2001. **265**: p. 888-894.
